# Majority of choice-related variability in perceptual decisions is present in early sensory cortex

**DOI:** 10.1101/207357

**Authors:** Michael J. Morais, Charles D. Michelson, Yuzhi Chen, Jonathan W. Pillow, Eyal Seidemann

## Abstract

While performing challenging perceptual tasks such as detecting a barely visible target, our perceptual reports vary across presentations of identical stimuli. This perceptual variability is presumably caused by neural variability in our brains. How much of the neural variability that correlates with the perceptual variability is present in the primary visual cortex (V1), the first cortical processing stage of visual information? To address this question, we recorded neural population responses from V1 using voltage-sensitive dye imaging while monkeys performed a challenging reaction-time visual detection task. We found that V1 population responses in the period leading to the decision correspond more closely to the monkey’s report than to the visual stimulus. These results, together with a simple computational model that allows one to quantify the captured choice-related variability, suggest that most of this variability is present in V1 as additive noise, and that areas downstream to V1 contain relatively little independent choice-related variability.

## Introduction

When we perform a challenging perceptual task, what fraction of the neural variability that relates to our variable perceptual decisions is present in early sensory cortex? One way to address this fundamental question is to measure neuronal activity in a relevant sensory cortical area during the temporal interval over which the decision is formed and assess the extent to which the measured activity is correlated with trial-to-trial variability in perceptual decisions. If such measurements are highly predictive of behavioral choice, then most choice-related variability must be present in this sensory area. This result would imply that other areas contain little choice-related variability that is independent of the variability present in the sensory area.

Previous studies of the relationship between neural and behavioral variability have discovered co-variations between responses of individual neurons in sensory cortex and a subject’s behavioral choices to repeated presentations of an identical near-threshold sensory stimulus (e.g. ***Britten et al., 1996***; ***Celebrini and Newsome, 1994***; ***Cook and Maunsell, 2002***; ***de Lafuente and Romo, 2005***; ***Dodd et al., 2001***; ***Nienborg and Cumming, 2006***, ****2009****; ***Palmer et al., 2007***; ***Purushothaman and Bradley, 2005***; ***Uka and DeAngelis, 2004***). These co-variations, however, tend to be very weak, implying that individual neurons capture only a small fraction of the neural variability related to decisions. The large amount of choice-related variability that is unaccounted for in single-neuron studies may be distributed across other (unmeasured) sensory neurons, and may also occur outside of the sensory area. Even if the majority of choice-related variability occurred in a distributed sensory cortical population, single neurons still may not provide strong predictions of choice because sensory neurons are only weakly correlated with one another (e.g. ***Zohary et al., 1994***), and therefore, one neuron’s responses are only weakly related to the population response. Thus, single-neuron studies cannot provide strong constraints on whether most decision-related neural variability is present in sensory cortex.

In contrast, methods for measuring neural population responses, such as voltage-sensitive dye imaging (VSDI) (***Grinvald and Hildesheim, 2004***), have the potential to provide stronger constraints by capturing a larger fraction of the choice-related variability within the sensory area. Because even simple, spatially localized stimuli evoke responses over a broad region of sensory cortex (***Chen et al., 2006***; ***Grinvald et al., 1994***; ***Palmer et al., 2012***), and because these sensory cortical neurons are only weakly correlated with one another, subjects can benefit from combining these distributed signals to inform decisions. Thus, perceptual decisions may be based on the combined activity from a large population of sensory neurons. In this case, population measures may show stronger co-variations with choice, and thus account for a greater fraction of the choice-related neural variability. However, such methods are likely to also contain various sources of choice-unrelated variability, such as choice-irrelevant neural variability and non-neural measurement noise, which can weaken the relationship between the measured neural activity and behavior.

To address the possibility that choice-related activity is distributed across many sensory neurons, we used VSDI to record from a large population of neurons in primary visual cortex (V1) of two monkeys while they performed a difficult visual detection task. In this task, the monkey reports the presence of a small low-contrast Gabor target that appeared on half of the trials by making a saccadic eye movement to target location as soon as it is detected (i.e., a reaction-time task). We placed the visual target so that it fell at the center of the visual area represented by the imaged neurons. Thus, we were able to capture most of the target-related signals that the monkey used to form decisions in our task. In other words, V1 provided the sensory evidence, and we eavesdropped on the relevant portion of V1 while the monkey formed a decision. The fast temporal dynamics of the voltage-sensitive dyes and our reaction-time task design allowed us to focus on choice-related signals during the brief period leading to the decision.

V1 signals during the pre-decisional period are variable, and this variability in the evidence is likely to affect the monkey’s choices and limit perception. This pre-decisional variability could arise from feedforward, recurrent and feedback sources. Our goal here is to quantify the total pre-decisional choice-related variability present in V1, irrespective of its source. Perceptual decisions are also likely to be affected by sources of variability downstream to V1 that are not accessible in V1 prior to the formation of the decision (e.g., decision noise, criterion fluctuations). Our primary goal here is to determine if the majority of the pre-decisional choice-related variability is present in V1. This is a fundamental open question.

To better understand the relationship between measurements of neural variability and perceptual decisions, we developed a simple computational model that allowed us to quantify the fraction of choice-related variability captured in a measurement of neural population responses independent of choice-unrelated variability. A second model allowed us to examine whether the features of the observed data are better-explained by additive noise or by a variable multiplicative gain. We use our results and models to determine a lower bound on the fraction of choice-related variability that occurs in V1 during performance of a detection task, and to characterize the structure of the choice-related variability in V1 population responses.

## Results

### Choice-related population activity in macaque V1

We trained two monkeys to perform a simple reaction-time visual detection task (Fig. 1). The monkey began each trial by directing gaze to a central fixation point. To obtain a reward, the monkey had to rapidly shift gaze to a small peripheral target during ‘target-present’ trials (50% of trials), or to maintain fixation on the central fixation point during ‘target-absent’ trials (the remaining trials). We used a small Gabor patch stimulus (sinusoidal grating in a 2-D Gaussian envelope) to effectively drive V1 cells, and presented the stimulus at one or two contrast levels near each monkey’s perceptual detection threshold to maintain task difficulty. Target-present trials were classified as ‘Hits’ if the monkey correctly detected the target and ‘Misses’ otherwise. Target-absent trials were classified as ‘False Alarms’ (FAs) if the monkey reported seeing a target and ‘Correct Rejects’ (CRs) otherwise (performances summarized in Appendix 1 Table 1).

**Figure 1.**
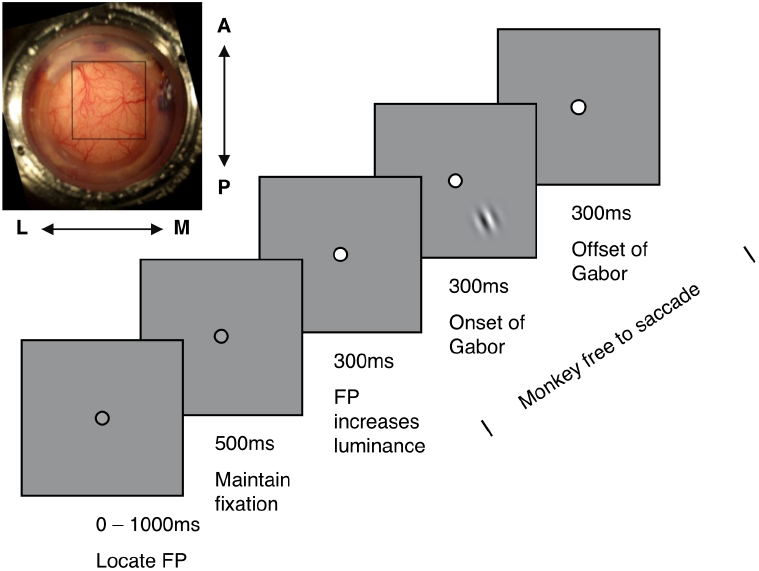
Visual detection task and simultaneous VSDI recording. *(Top left)* Cranial window over dorsal V1. Black square indicates VSDI imaging area of 10 x 10 mm^2^ in the left hemisphere. Letters indicate approximate cortex orientation (A – anterior; P – posterior; M – medial; L – lateral). *(Center)* Behavioral task. Each trial begins when the subject locates and fixates a central fixation point. After a 500 ms delay during which the subject is required to maintain fixation, the fixation point undergoes a slight increase in luminance to indicate the start of the task portion of the trial. Following a 300 ms delay, a Gabor stimulus appears on 50% of trials. The subject is required to make a saccadic eye movement to the stimulus location when it appears, or to maintain fixation on the central fixation point for an additional 900 ms when it does not appear. The subject receives a drop of juice or water following correct choices, and no reward following incorrect choices.

While the monkeys performed the task, we used voltage-sensitive dye imaging (VSDI) to measure population responses from V1, an area likely to play a central role in performance of such tasks. VSDI measures changes in membrane potential from large populations of neurons with a dominant contribution from layer 2/3 (***Chen and Seidemann, 2012***; ***Petersen et al., 2003***). Previous work showed that this technique can be sufficiently sensitive to outperform monkeys in a similar detection task when combining single-trial VSDI signals optimally (***Chen et al., 2006***, ***2008***). Furthermore, because V1 contains a topographic map of visual space, the target activates a limited region in V1 (***Chen et al., 2006***; ***Sit et al., 2009***), and this region can be precisely localized and entirely captured by VSDI. Thus, VSDI may be capable of capturing a large portion of choice-related variability in V1.

What are the expected trial-to-trial co-variations between VSDI responses and the monkey’s choices? Consider a scenario in which all choice-related variability is present in V1 during a detection task. Intuitively, in this case, FA responses should resemble the average target-evoked responses, with amplitudes that exceed the internal detection criterion, even though the target is absent. Similarly, Miss responses should fall below the internal detection criterion, even though the target is present. Thus, in this case, we expect the average V1 responses on FAs to exceed responses on Misses. On the other hand, if most of the choice-related variability occurs downstream to V1, V1 responses should be only weakly correlated with choice, and the average responses in FAs should be much weaker than in Misses.

Results from a representative VSDI experiment (Fig. 2A) reveal large choice-related modulations at the level of neural populations in V1. On average, the stimulus evoked a broad Gaussian-shaped response across the population of neurons, consistent with previous results (***Chen et al., 2006***, ***2008***; ***Chen and Seidemann, 2012***; ***Sit et al., 2009***). The mean stimulus-present and -absent maps were each further partitioned into mean Hit and Miss maps, and mean FA and CR maps, respectively. These maps reveal robust choice-related differences. Similar choice-related activity is also observed in the grand average results combined across all of our data sets (Fig. 2B; see Fig. 2 – Supplementary Fig. 1 for results from individual monkeys). These maps show that choice-related activity, as revealed by the Hit-Miss and FA-CR difference maps, is distributed broadly across the neuronal population rather than being restricted to a small region. To quantify these choice-related differences, we computed the time- and trial-averaged amplitudes of the VSDI response for each trial outcome (Fig. 2C). Average amplitudes are significantly larger in Hits than in Misses (p<0.001, bootstrap test), significantly larger in FAs than in CRs (p<0.001, bootstrap test), and significantly larger in FAs than in Misses (p<0.01), suggesting that in this task, the population activity in V1 corresponds more closely to the subject’s choice than to the stimulus.

**Figure 2.**
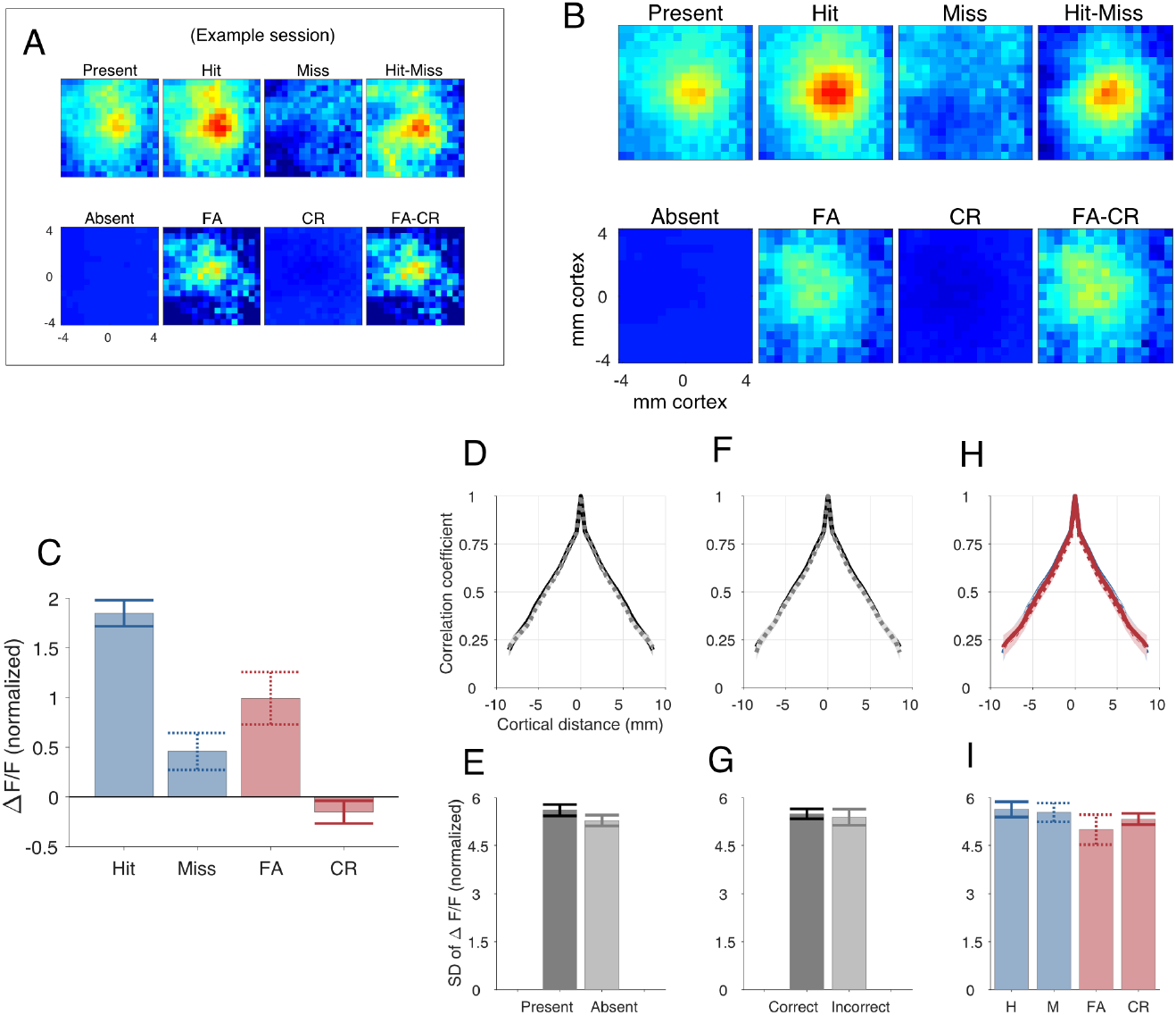
Choice-related population activity in V1. **A**, Example VSDI experiment in Monkey T. Average cortical activity maps separated by stimulus condition (rows) and behavioral choice (columns). Upper row of panel A represents neural activity during target-present trials; lower row shows activity during target-absent trials. Activity maps are shown separately for ‘choose target-present’ (Hit and FA) and ‘choose target-absent’ (Miss and CR) trials. ‘Hit-Miss’, difference map of activity during Hit and Miss trials. ‘FA-CR’, difference map of activity during FA and CR trials. Cortical maps represent activity averaged over a short time interval after stimulus onset (see Methods). **B-C**, Aggregate summary of VSDI results collapsed over 22 experiments in Monkey T and 5 experiments in Monkey C. **B**, Average cortical activity maps separated by stimulus condition (rows) and behavioral choice (columns), with the same organization as panel A. **C**, Bar plot showing the mean and SE (bootstrapped) of response amplitudes (computed over 2.5×2.5 mm^2^ centered at the peak response) for each of the 4 behavioral categories. Means are weighted averages, with weights proportional to the SNR in each experiment. Bar plots were normalized for display such that the mean target-absent response was 0 and the mean target-present response was 1. **D,F,H**, Spatial correlation coefficients computed across all pairs of pixels, plotted as a function of cortical distance; **E,G,I**, standard deviations of normalized response amplitudes. Error bars and shaded areas indicated 95% con1dence intervals (bootstrapped). The noise levels and spatial correlation profiles were statistically indistinguishable during target-present vs. target-absent trials (**D-E**), during correct vs. incorrect choices (**F-G**), and during each of the four behavioral outcomes (**H-I**). **Figure 2–Figure supplement 1.** Choice-related population activity in V1, per animal.

These choice-related differences in the means of VSDI activity are not reflected in the spatial correlation profiles of VSDI activity (Fig. 2D, F, H), nor in the standard deviations of that activity (Fig. 2E, G, I). We compare Present vs. Absent trials (Fig. 2D, E), Correct vs. Incorrect trials (Fig. 2F, G), and Hit vs. Miss vs. FA vs. CR trials (Fig. 2H, I), and no pairwise tests yield significant differences (for SDs, t-tests between groups, all p>0.05; for spatial correlations, bootstrap tests shuffling groups, all p>0.05). These null results agree with prior work which demonstrated that variability in VSDI measurements of V1 population responses is consistent with Gaussian additive noise (***Chen et al. 2006***, discussed further below).

To further examine the co-variation between V1 population responses and behavior, we analyzed the relationship between reaction times and time-averaged VSDI activity. This analysis reveals a negative correlation between the two for Hit trials; that is, higher-than-average V1 responses co-occur with lower-than-average reaction times (Fig. 3, p<0.005). The relationship is similar on FA trials, but is not significant (p=0.156), perhaps in part due to the lower trial counts.

**Figure 3.**
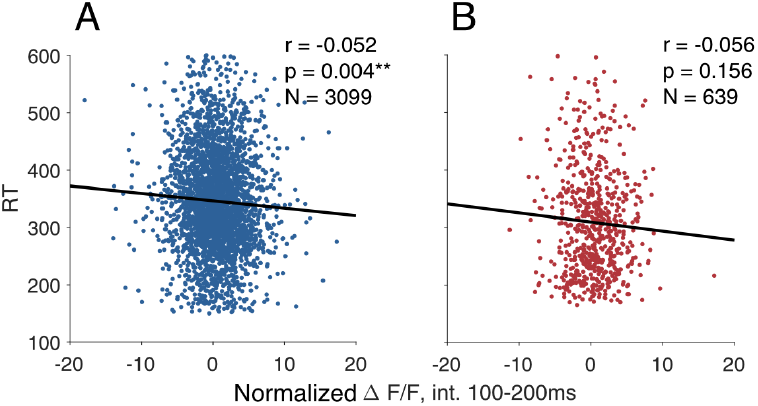
Correlation between reaction time and VSDI activity. Reaction time is negatively correlated with VSDI activity during the 100ms window from 100-200ms post-stimulus onset, significantly for Hit trials (**A**) and not for FA trials (**B**).

Additional analysis of eye position suggests that our results are not due to small variations in eye movements (Appendix 1 Table 2, see Methods and Materials for details).

### Computational model for assessing the fraction of choice-related variability in V1

Our results show significant choice-related variability in V1 but cannot, by themselves, determine whether most of the choice related variability is present in V1. To address this question, we developed a simple computational model of the pooled single-trial VSDI responses and used this model to obtain a quantitative estimate of the fraction of choice-related variability present in V1. In our model, the pooled sensory signal in target-absent and target-present trials is assumed to be normally distributed with variance 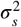 and means zero and one, respectively (Fig. 4, ‘S’). During formation of the decision, additional independent decision-related Gaussian noise with variance 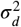 is added to the pooled sensory response (Fig. 4, ‘D’). The total decision variability, which controls the subject’s accuracy, is thus the sum of the contributions from these two independent sources of decision-related variability, 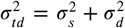. Note that this formulation does not distinguish among the various possible sources of the pre-decisional choice-related V1 variability. Thus, 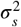 could have bottom-up (feedforward) and top-down (feedback) contributions, while 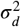 captures choice-related variability outside of V1 that is independent of V1 variability and is therefore inaccessible from V1 measurements.

**Figure 4.**
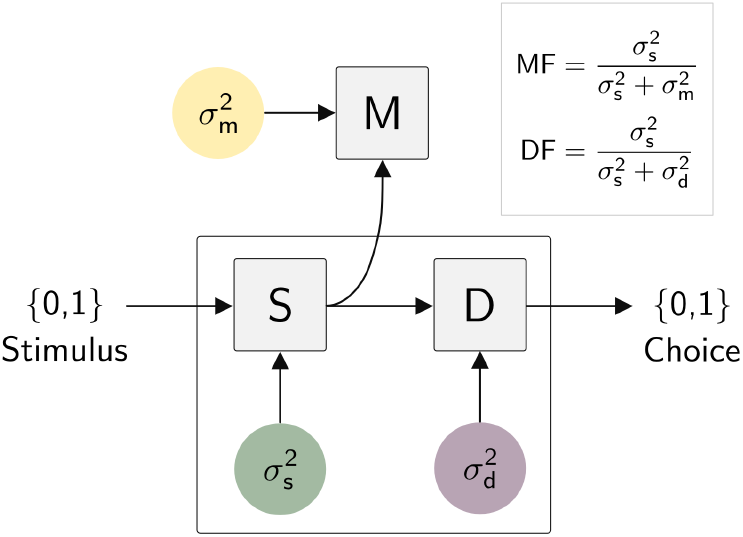
Theoretical framework. Model of a subject’s formation of a binary decision and a researcher’s measurement of neuronal responses. Stimulus-induced responses in an early sensory area come from distributions with means 0 and 1 and variance 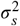 (green). Zero-mean noise with variance 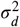 representing downstream variability (purple) is added to form the decision variable. The subject compares the decision variable to a criterion to form a decision on each trial. The upper portion of this panel represents stages in a researcher’s measurement, in which zero-mean choice-unrelated noise with variance 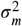 (yellow) is added to the sensory representation to form the measured signals. The measured signals are then analyzed with respect to the subject’s binary choices on a trial-by-trial basis.

The subject reports ‘target present’ if the decision variable exceeds a criterion, and ‘target absent’ otherwise. As experimenters, we have access to the sensory neural responses (Fig. 4, ‘S’), but we assume that our measurements are mixed with independent, choice-unrelated noise with variance 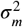 (Fig. 4, ‘M’) which combines neural and non-neural sources of variability. The total variability in the signal thus includes both the choice-related sensory variability and the choice-unrelated measured variability, 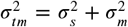. Here we employ a slightly richer model that takes into account multiple target contrasts and experiment-to-experiment variations in VSDI signal quality (see Methods), and can confirm that the model’s assumption of equivariant Gaussian responses for H, M, FA, and CR trials is an accurate description of our data (see Appendix 1 Fig. 1).

Our model can be used to estimate the fraction of the total decision-related variability that is present in the sensory area, 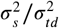, which we abbreviate DF (‘decision fraction’). A second quantity of interest is the fraction of the total measured variability that is choice-related, 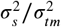, which we abbreviate MF (‘measured fraction’). Consistent with our intuition, the model’s behavior shows that the difference between the mean response in FAs and Misses is directly related to DF. When DF is high, the mean FA exceeds the mean Miss (Fig. 5A, dashed red vs. dashed blue arrows). When DF is low, the mean Miss exceeds the mean FA (Fig. 5C), and the two means are equal at some intermediate value of DF. In fact, there is a linear relationship between the difference in these means and DF (Fig. 5E, black curve). In addition, as DF decreases, the correlation between the measured response and choice drops. A common metric for measuring this correlation is choice probability (CP), which measures the discriminability of two distributions of neuronal responses that belong to the same stimulus condition but to two different behavioral choices, such as ‘Hits’ (solid blue line) and ‘Misses’ (dashed blue line). As DF decreases, CP drops from a value of 1 (perfect discriminability) toward 0.5 (no discriminability) (Fig. 5E, gray curve). Importantly, while CP is strongly affected by measured variability that is choice unrelated (Fig. 5D,F), such variability has no effect on the means of the responses tied to each of the 4 behavioral outcomes. The invariance of the means with choice-unrelated variability allows our model to estimate DF independent of MF.

**Figure 5.**
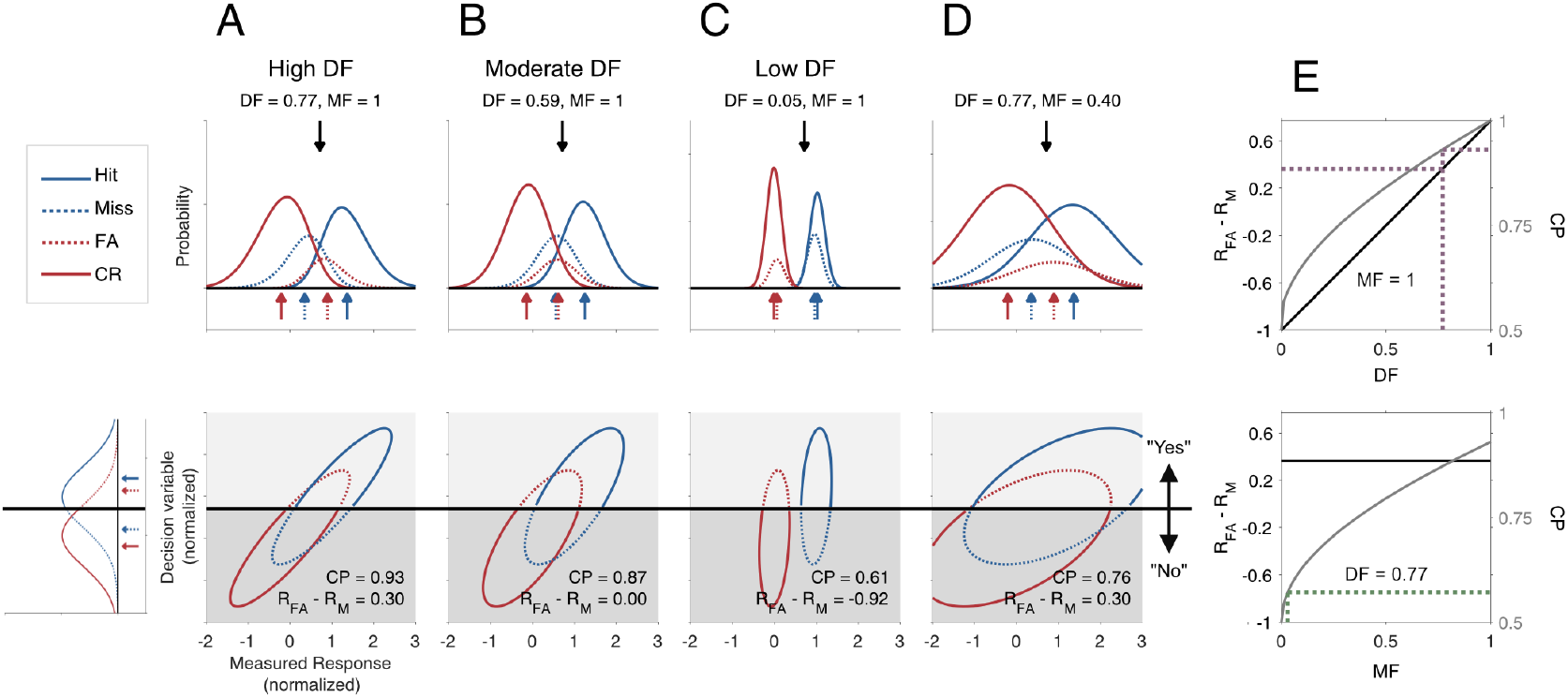
Expected relationship between observable quantities and the underlying properties of a decision. **A-D**(*lower*), Joint distributions of the decision variable (ordinates) and model-predicted pooled neuronal responses (abscissae) under 4 noise scenarios, ranging from 77% of choice-related variability occurring in a measured brain area (**A**) to only 5% of choice-related variability captured by the measurement and the remaining 95% occurring downstream (**C**). In panels **A-D**, marginal distributions are shown for the four behavioral categories. The marginal distributions of the ordinate of panel **A** represent the decision variable which is controlled by the animal’s overall accuracy and is therefore fixed in all panels. Red and blue arrows indicate mean neuronal response amplitude for each of the 4 behavioral categories. In panel **B**, the mean neuronal response amplitude for ‘Miss’ trials is identical to that of FA trials (dotted red arrow). Black arrows (top panels) and horizontal black lines (bottom panels) indicate the decision criterion. Mean response in FA minus mean response in Miss (*R*_*FA*_ − *R*_*M*_), and choice probability (CP) values (see text for details) are reported for each scenario. In panels **A-C**, no choice-unrelated variability was added to the expected measured responses (i.e., MF = 1). **D**, Same as panel **A**, after adding a large amount of choice-unrelated variability. **E**, Relationship between *R*_*FA*_ − *R*_*M*_ (left ordinate, black), CP (right ordinate, gray), and DF in the simplified model. Dotted blue lines indicate DF solution from the full model applied to the real data for the choice-triggered pooling rule in the absence of choice-unrelated noise (i.e., MF=1) and corresponding *R*_*FA*_ − *R*_*M*_ and CP values in the simplified model. F, The relationship between *R*_*FA*_ − *R*_*M*_ (left ordinate, black), CP (right ordinate, gray), and MF in the simplified model when DF = 0.77. While *R*_*FA*_ − *R*_*M*_ is independent of MF, CP varies systematically with MF. Dotted blue lines indicate MF solution from the full model applied to the real data for the choice-triggered pooling rule (MF = 0.03) and corresponding CP value in the simplified model. The large amount of choice-unrelated variability reflects a combination of neural variability (e.g., independent neural variability that is captured by our measurements but does not affect choice) and non-neural measurement noise.

By applying our model to our results, we found that the majority of choice-related variability is present in V1. To estimate the fraction of choice-related variability captured by our VSDI measurements, we combined the signals over space and time into a single value for each trial, and examined the distribution of these scalar responses across trials. We first consider results under a simple pooling rule that uses weights proportional to the average choice-triggered responses (see Methods). The mean pooled signal is significantly higher in FA than in Miss trials, consistent with Fig. 2C. According to our model, these means imply that DF = 0.77 (95% CI: [0.56 1.0]), or in other words, 77% of the choice-related variability is present in V1. Similar results were obtained with a wide range of plausible candidate spatial pooling rules (Table 1), providing evidence that in our task, the majority of choice related activity is present in V1.

**Table 1.** Model solutions for 9 alternative pooling rules. We repeated our maximum likelihood estimation procedure (see Methods and Materials) for each of 9 alternative candidate pooling rules to obtain estimates for DF and MF. DF estimates ranged from 0.72 to 0.99, and MF estimate ranged from 0.03 to 0.07 across pooling rules. For each pooling rule, we also computed choice probability expected under the simplified model associated with the DF and MF solutions. Values in parentheses indicate 95% bootstrapped confidence intervals. ‘CP (MF = 1)’ refers to the choice probability value associated with each pooling rule’s DF solution, in the absence of choice-unrelated noise (i.e., MF = 1). ‘CP’ refers to the choice probability value associated with each pooling rule’s DF solution, after the measured neural signals have been corrupted by the addition of choice-unrelated noise consistent with that pooling rule’s MF solution. ‘NS (MF = 1)’ and ‘NS’ refer to the neural sensitivity, reported analogously to CP.

| Pooling Model | Decision Fraction (DF) | Measurement Fraction (MF) | CP (MF = 1) | CP | NS (MF = 1) | NS |
| --- | --- | --- | --- | --- | --- | --- |
| 1 - Fitted Choice-triggered Average | 0.77 (0.40, 1.00) | 0.03 (0.01, 0.04) | 0.93 | 0.56 | 0.90 | 0.58 |
| 2 - Fitted Average Evoked | 0.77 (0.42, 1.00) | 0.03 (0.01, 0.05) | 0.93 | 0.56 | 0.90 | 0.58 |
| 3 - Gaussian (sigma = 0.5mm) | 0.97 (0.70, 1.00) | 0.07 (0.04, 0.09) | 0.99 | 0.61 | 0.88 | 0.61 |
| 4 - Gaussian (sigma = 1.0mm) | 0.96 (0.67, 1.00) | 0.06 (0.03, 0.07) | 0.99 | 0.60 | 0.88 | 0.60 |
| 5 - Gaussian (sigma = 2.0mm) | 0.86 (0.53, 1.00) | 0.04 (0.02, 0.06) | 0.96 | 0.58 | 0.89 | 0.59 |
| 6 - Gaussian (sigma = 3.0mm) | 0.77 (0.40, 1.00) | 0.03 (0.01, 0.05) | 0.93 | 0.56 | 0.90 | 0.58 |
| 7 - Gaussian (sigma = 4.0mm) | 0.72 (0.36, 1.00) | 0.03 (0.01, 0.04) | 0.92 | 0.56 | 0.91 | 0.58 |
| 8 - Optimal (Present vs. Absent) | 0.98 (0.65, 1.00) | 0.05 (0.02, 0.07) | 0.99 | 0.59 | 0.88 | 0.59 |
| 9 - Optimal ("Yes" vs. "No") | 0.99 (0.68, 1.00) | 0.05 (0.03, 0.07) | 0.99 | 0.60 | 0.88 | 0.60 |

Suppose we operationalized the spatial pooling rules in Table 1 as decoders based on the VSDI signals. While high DF alone implies that V1 would have strong decoding performance, the combination with low MF implies that this would-be strong performance is corrupted by substantial choice-unrelated noise. The parametrization of variability in our model allows us to disambiguate these competing influences, and we can “remove” the decision-unrelated measured variability (fix 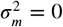 such that MF = 1) to observe the uncorrupted decoding performance when DF is high. We observe modest CP = 0.56 (‘CP’), as the total variability in V1 is dominated by decision-unrelated variability (low MF). In the absence of this decision-unrelated variability, though, we would observe CP = 0.93 (‘CP (MF = 1)’). We also measured our ability to decode the visual stimulus from the single-trial pooled V1 responses – defined as neural sensitivity (NS). The choice-unrelated variability similarly affects NS. Similar to our results with CP, we observe NS = 0.58 (‘NS’), and NS = 0.90 in the absence of decision-unrelated variability (‘NS (MF=1)’)(see Table 1 for results under other spatial pooling rules).

### Effect of partial overlap between subject and researcher’s sensory neural pools

Our model (Fig. 4) assumes that the subject and researcher share equal access to the V1 population. However, population measures such as VSDI are limited to finite spatial extent, spatial resolution, and cortical depth. Thus the subject and researcher’s pooled sensory responses may overlap only partially. To explore the possible impact of partial overlap, we extended our model by simulating V1 as a 2-dimensional patch of neurons/pixels with signal and noise characteristics that matched those from the real data (see Supplemental Methods). We then performed simulated experiments in which the researcher, the subject, or both, are given independent access only to a random sub-population of the simulated cortex. Our results (Fig. 6) show that the DF estimated by our model is a lower bound on the true fraction of choice-related variability present in sensory cortex, because some of what our model considers as downstream noise may actually reside in V1.

**Figure 6.**
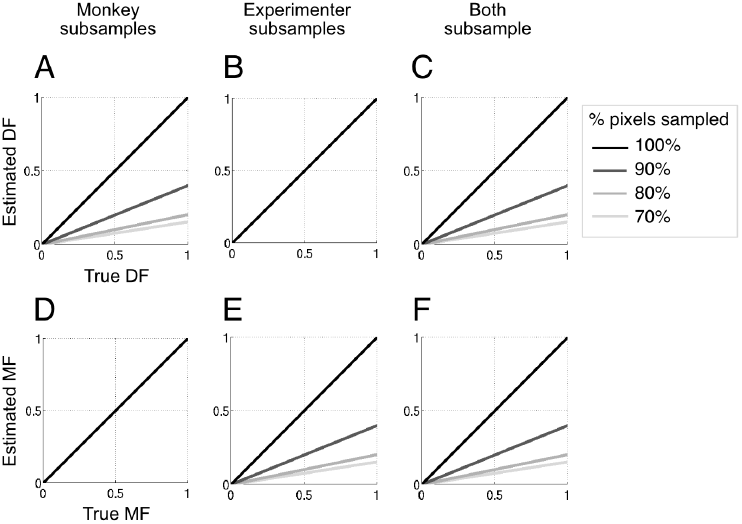
Sub-sampling analysis. Relationships between DF and MF values inferred using the maximum likelihood method (ordinates) and those used to generate the simulation (abscissae) during conditions in which the simulated monkey sub-samples the cortical population (**A**,**D**), the simulated experimenter sub-samples (**B**, **E**), and both the simulated monkey and experimenter sub-sample (**C**, **F**) (see Methods and Materials). Black lines correspond to the case in which our initial assumption of equal access is met (see text for details). Grey lines indicate the percentage of pixels sampled. This figure demonstrates that the DF and MF values inferred with the maximum likelihood method provide lower bounds on their true values.

### Computational model for assessing additive versus multiplicative variability in V1

We supplemented the scalar likelihood model with a targeted series of qualitative simulations, to investigate whether the choice-related variability we observe in our data was consistent with additive noise, as we assumed in our pooled scalar likelihood model, or whether those results can emerge from multiplicative gains. We generalized the scalar-valued likelihood model (cf. Fig. 4) to a pair of matrix-valued models that directly simulates VSDI activity maps at the pixel resolution of our data, and inherits the partitioning of choice-related sensory noise, decision noises and choice-unrelated measured noise, (see Methods and Materials and Appendix 2 for more detailed derivations). We specifically investigated the shared sensory noise; the measured and decision noises remained additive. In the additive noise model, the sensory noise is spatially correlated Gaussian noise *added* to the stimulus, and in the multiplicative gain model, the sensory noise is substituted for spatially correlated Gamma gains *multiplied* to the stimulus. Each model is optimized to recapitulate the key features observed of our VSDI data – the subject’s performances in terms of Hits, Misses, FAs, and CRs, as well as trial-averaged activity maps and their spatial correlations profiles (cf. Fig 2) – under these different sources of variability.

Both additive signal noise and stochastic multiplicative signal gain models can readily match the observed proportions of Hits, Misses, FAs, and CRs (Fig. 7A,F) as well as the spatial correlation profiles (Fig. 7D,I) and response variability (Fig. 7E,J). However, while the additive noise model matches also the empirical VSDI pooled response maps of all trial types (Fig. 7B,C, and Fig. 7G,H, *resp.*), the multiplicative noise model generates FAs differently – namely that FAs do not present with increased VSDI signal relative to Misses and CRs as they do in the empirical data (Fig. 7G,H).

**Figure 7.**
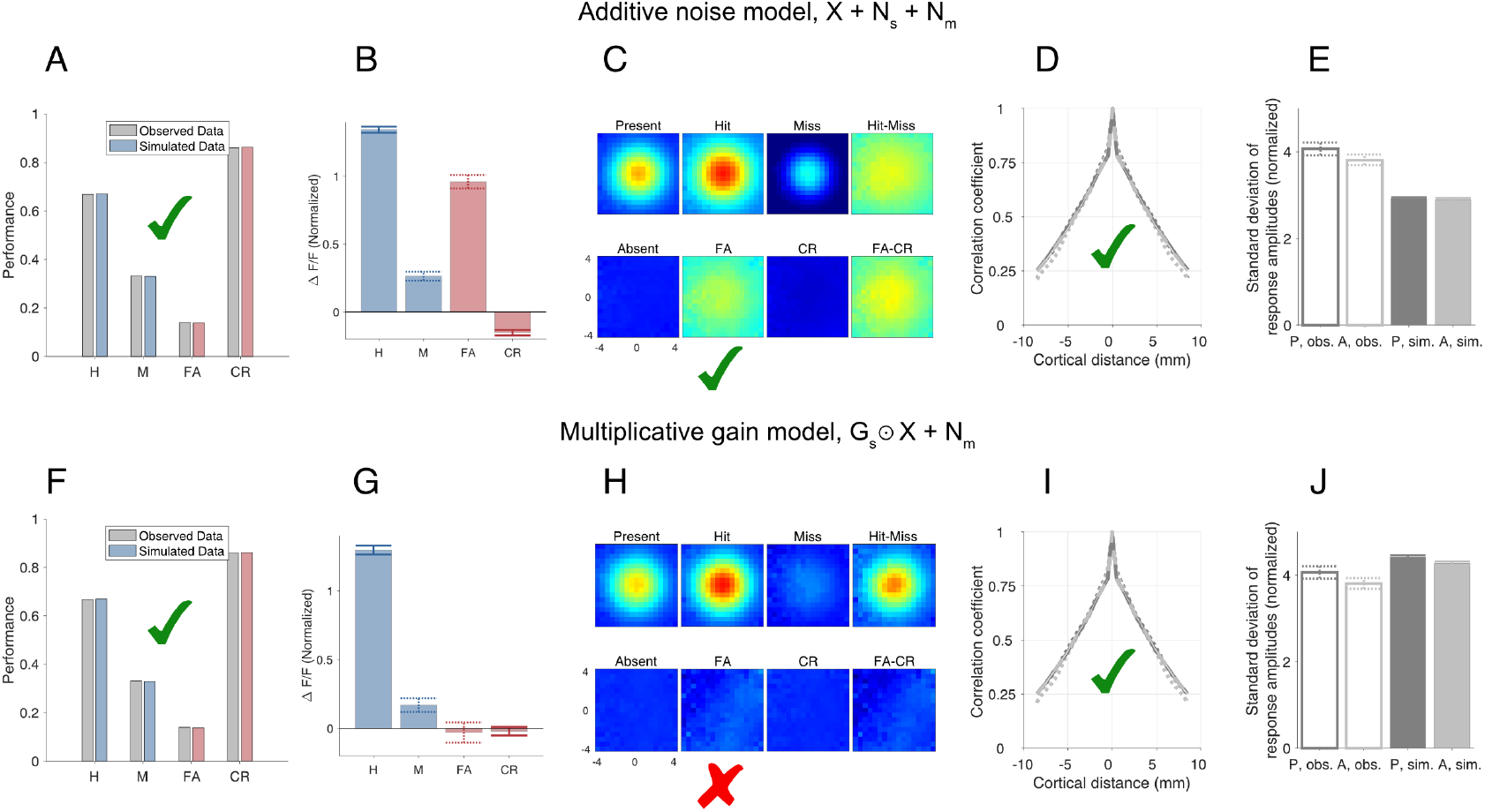
Performance, mean VSDI activity maps, and spatial correlations under different simulated sources of shared variability. (**A-E**) Additive noise model in which the observed VSDI activity is generated by a stimulus profile (either on = X, or off = 0), plus spatially correlated Gaussian-distributed noise. All key features of the data (performance, spatial correlations, VSDI activity maps with *R*_*FA*_ > *R*_*M*_) are replicated by this model. (**A**) Performance for Hit, Miss, False Alarm, and Correct Reject trials in observed dataset (gray) and simulated dataset (Blue & Red). (**B**) Mean and SE (bootstrapped) of response amplitudes for Hit, Miss, False Alarm, and Correct Reject trials in simulated dataset; cf. Fig. 2C (**C**) Mean VSDI activity maps by trial outcome in a simulated 8×8 mm^2^ area; cf. Fig. 2A. (**D**) Spatial correlations and (**E**) pooled response standard deviations for observed present and absent trials (dotted black and gray lines, resp., and outlined black and gray bars) and simulated present and absent trials (solid black and gray lines, resp., and solid black and gray bars), each with 95% CIs; cf. Fig. 2D,E. (**F-J**) Multiplicative gain model in which the activity is generated by the stimulus profile, times a Gamma-distributed gain. Some features of the data (performance, spatial correlations) are replicated by this model, but in the VSDI activity maps, *R*_*FA*_ is 0. Panels organized as in Panels (**A-E**).

Because a stochastic gain *G* multiplying a signal *X* contributes zero variability when *X* = 0 (i.e., in target-absent trials), FAs can emerge in a multiplicative gain model only from the downstream decision-related variability captured by 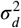. Therefore, no elevated VSDI activation is observable on FA trials in this case, inconsistent with our results (Fig. 2). Together, these simulations further support that choice-related sensory variability in these VSDI signals is consistent with Gaussian additive noise and inconsistent with stochastic multiplicative signal gain.

### Dynamics of choice-related variability in V1

We next examined the dynamics of the choice-related variability in V1. In order to perform the detection task, the monkey must form a decision by accumulating weak and noisy evidence provided by populations of V1 neurons. Therefore, if most of the choice-related variability that we observe in V1 is post-decisional, we would expect the onset of the choice-related V1 signal to be significantly delayed relative to the onset of the target-evoked V1 response. If, on the other hand, the choice-related signal reflects V1 variability that is part of the evidence that contributes to the decision, we would expect the dynamics of the choice-related signal to be similar to the dynamics of the target-evoked response. Our next step was to compare the dynamics of the choice-related signal with the dynamics of the target-evoked response.

Our results reveal tight coupling between the dynamics of the choice-related and target-evoked population responses in V1, providing evidence against a major post-decisional source for the observed choice-related variability (Fig. 8). Because the target is small and has a very low contrast, the target-evoked response in V1 – mean target-present minus mean target-absent responses – has a delayed onset and a slow time-to-peak (solid gray curves, Fig. 8), consistent with previous studies (***Sit et al., 2009***). If we separate all target-present trials based on choice, we find that the separation between Hit and Miss responses (solid and dashed blue curves, resp., Fig. 8A) starts at approximately the same time as the target-evoked response. Similarly, if we separate all target-absent trials based on choice, we find that the separation between FA and CR responses (dashed and solid red curves, resp., Fig. 8B) starts at approximately at the same time as the target-evoked response. The horizontal bars at the bottom of both panels of Fig. 8 denote time windows of significant (p<0.05) target-evoked (gray) and choice-related activities (H-vs-M, blue, and FA-vs-CR, red; see Fig. 8 – Supplementary Fig. 1 for 95% CIs). Their errorbars are 95% CI on the *onset* time, and none are significantly different. Therefore, the dynamics of the choice-related response in V1 are inconsistent with a dominant post-decisional source to the observed choice-related variability.

**Figure 8.**
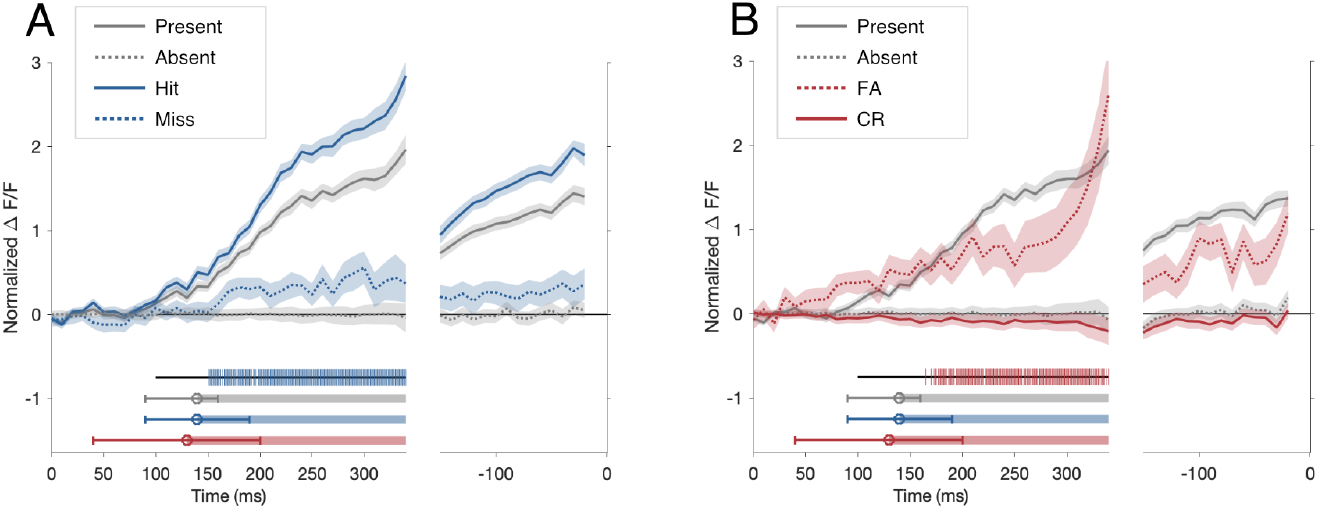
Temporal dynamics of choice-related activity. Time evolution of mean normalized response amplitudes and choice-related differences for each of the 4 behavioral outcomes. Shaded areas indicate bootstrapped standard errors. Hash marks at the bottom of the panels indicate saccadic onset times of the respective trial types. Horizontal bars at the bottom of the panels indicate significant choice-related differences (for bootstrapped CIs used in statistical testing, see associated Supplementary Fig. 1). Errorbars are bootstrapped CIs atop the onset time of these differences. Traces for trials in which the monkey did not make an eye movement (Miss and CR trials) are shown for intervals re-sampled from those of the Hit and FA trials. Underneath both panels, horizontal bars reflect significant choice-related activity that onsets at similar times across all trial types. Black horizontal line – averaging period for quantitative analysis (see Methods). **A**, Temporal dynamics on target-present trials; target-present and -absent responses are shown in gray (solid and dashed, resp.). The gray horizontal barsshow the temporal window in which their difference (mean ‘P’ minus mean ‘A’) is significantly discriminable from zero; errorbars are a 95% CI on the onset time. Hit trials (solid blue) accumulate activation over the course of the trial, while Miss trials (dashed blue) do not. The blue horizontal bar shows the discriminability of their difference over time (mean ‘H’ minus mean ‘M’). **B**, Temporal dynamics on target-absent trials; same as panel **A** but for FA trials (dashed red) and CR trials (solid red), respectively. The red horizontal bar shows the discriminability of their difference over time (mean ‘FA’ minus mean ‘CR’). **Figure 8–Figure supplement 1.** 95% Confidence regions for significant choice-related differences

The observed dynamics of the choice-related signal in V1 (Fig. 8) are also inconsistent with another possible source of choice-related variability – fluctuations in top-down attentional or preparatory signals. We previously discovered such attentional and preparatory signals in population responses in V1 while monkeys performed a similar detection task (***Chen and Seidemann, 2012***). Because in our detection task the timing of target onset was cued in advance, the attentional and preparatory signals in V1 were anticipatory in nature and started well before stimulus onset (***Chen and Seidemann, 2012***). Therefore, if fluctuations in these anticipatory signals were a major source of choice-related variability, we would expect choice-related signal to emerge well before stimulus onset. Our finding that the choice-related V1 signal only occurs around the time of the target-evoked response (Fig. 8) is therefore inconsistent with this possibility.

## Discussion

Here we present two main findings. First, in a visual detection task, there is strong choice-related activity in V1 that is distributed across tens of mm^2^ of cortex. This finding suggests that subjects combine sensory signals from a broad region of cortical space in order to inform decisions. Second, V1 population responses correspond more closely to the subject’s choices than to the stimulus. To estimate the fraction of choice-related variability present in V1, we developed a simple model for studying the relationship between choice-related activity and noisy measures of neural population responses. Applying this model to our data, we find that V1 population responses account for the majority of the subjects’ choice-related neural variability. Our results are consistent with previous studies that used indirect methods to infer the sources of behavioral variability (***Brunton et al., 2013***; ***Osborne et al., 2005***), and with recent studies arguing that correlated noise present in early sensory areas is the primary factor limiting perception (***Moreno-Bote et al., 2014***; ***Pitkow et al., 2015***).

The choice-related signals we observed are likely to contribute to the decision and limit behavior because they co-occur in space (Fig. 2) and in time (Fig. 8) with the V1 signals that the subject must use in order to perform the task. In other words, to perform our detection task, monkeys must rely on signals that pass through the imaged area in V1 during a brief pre-decisional temporal interval. These pre-decisional V1 signals are variable, and this variability must therefore affect the monkeys’ decision. The sources of the V1 variability that affect decisions are not known, and are likely to include feedforward and feedback contributions. While feedback contributions to choice-related activity are well documented (e.g. ***Nienborg and Cumming, 2009***), most of the previously observed feedback contribution to choice-related signals may be post-decisional, and the relative contribution of feedback to the variability in the sensory evidence that is used to form the decision is unknown. Our study focuses on a separate open question – what is the relative contribution variability in V1 vs. independent variability downstream to V1 (e.g., decision or criterion variability) to the variability that limits perception in our detection task.

Overall perceptual sensitivity can be limited by three factors: the quality of the sensory evidence in early sensory cortex, the efficiency of the decoding mechanisms that convert these sensory signals into a perceptual decision, and the magnitude of downstream decision noise (***Seidemann and Geisler, 2018***). Our results suggest that in our task, the contribution of sensory variability to variability in perceptual decisions is larger than the contribution of decision noise. However, our results do not speak directly to the relative contribution of inefficient decoding or pooling to limiting perceptual sensitivity. In our previous work, we have shown that the sensitivity of V1 population responses exceeds the behavioral sensitivity of the subject in detection tasks even when considering only the locally pooled neural response (i.e., the coarse population response at the retinotopic scale). These results imply that decoding or pooling inefficiencies play an important role in limiting perceptual sensitivity. Recent experiments measuring neural sensitivity of populations of thousands of individual neurons in mouse V1 to stimulus orientation showed that neural sensitivity is orders of magnitude higher than mouse behavioral sensitivity (***Stringer et al., 2021***). These results are consistent with the hypothesis that decoding or pooling inefficiency is a major factor limiting perceptual sensitivity, though the degree of this inefficiency is likely to be species and task dependent (see ***Seidemann and Geisler 2018*** for a related review).

Our results demonstrate that most of the variability that limits performance in our detection task is present in V1. Moreover, a second computational model (Fig. 7) suggests that the source of the choice-related variability observed in the VSDI data is additive sensory noise rather than variable multiplicative gain. Specifically, variable multiplicative gain cannot yield population activity in which the mean FA response exceeds the mean Miss response, one of our key observations (Fig. 2).

What might be the sources of the pre-decisional V1 variability? Because the stimulus sequence in our task was random, top-down signals that add a variable bias to V1 activity (e.g., signals reflecting trial-by-trial expectations) could only be detrimental to task performance. Since our results were obtained from highly trained subjects, the presence of such detrimental top-down signals in our results is likely to have been minimized. The choice-related signals reported here are selective to the location of the target (see Hit-Miss and FA-CR maps in Fig. 2). As such, they are unlikely to reflect additive fluctuations in arousal/alertness. The choice-related signals reported here are selective to the location of the target (see Hit-Miss and FA-CR maps in Fig. 2). As such, they are unlikely to reflect additive fluctuations in arousal. Another potential source of top-down variability in V1 is fluctuations in spatial attention. However, in a previous study of spatial attention in V1 which employed a similar detection task with similar temporal cueing (a change in fixation point brightness 300 ms before the time of target onset), we observed attentional effects that precede target onset and, while being spatially selective, spread over a much larger spatial extent than the target-evoked responses (***Chen and Seidemann, 2012***) and the choice-related signals reported here. Thus, these top-down attentional signals appear to have different spatiotemporal dynamics from the choice-related response observed here (Fig. 2D; Fig. 8). Finally, attention and alertness have been reported to have a multiplicative effect on sensory responses (e.g. ***Reynolds and Heeger, 2009***). Such multiplicative effects cannot account for the signals that we observed in FA trials in which no sensory stimulus was present (Fig. 7). These considerations suggest that in our task, trial-by-trial fluctuations in attention and alertness may have a relatively small contribution to pre-decisional choice-related variability in V1. Fluctuations in attention and alertness might instead contribute primarily to V1 variability that the decoder is insensitive to (i.e., variability that falls in the decoder’s null space), and is therefore choice-unrelated.

Regardless of its origin, we show here that in a detection task, the majority of choice-related variability can be found in V1 during the epoch leading to the decision, and that this variability has spatiotemporal dynamics similar to those of the target-evoked response. These results provide strong constraints on models that link neural responses in early sensory cortex and perceptual decisions.

Our result that the sensory population signals correspond more closely to the subject’s choices than to the stimulus is similar to fMRI findings reported previously in human V1 (***Ress and Heeger, 2003***). However, due to the sluggish nature of fMRI signals and the delayed-response task design, the previous study was unable to determine whether the choice-related activity observed in sensory cortex occurred before or after the subject’s decision. The previous study found similar activity in Hits and FAs and similar activity in Misses and CRs, suggesting that the previous results were dominated by activity occurring in an epoch in which stimulus-related signals had already vanished. In contrast to the slow and indirect nature of the fMRI signal, the fast temporal dynamics of the voltage-sensitive dyes and our reaction-time task design allowed us to focus on choice-related signal that occur during the formation of the decision.

Our study speaks to fundamental questions about the relationship between noisy sensory representations and perceptual decisions. Here we show that decision-related neural activity is distributed broadly in early sensory cortex, suggesting that subjects pool information over a large population of sensory neurons to inform decisions, rather than relying on a small group of highly informative neurons. Second, we provide evidence that most choice-related neural variability is already present in early sensory cortex prior to the decision, suggesting that downstream circuits add little independent variability that affects decisions. An important goal for future studies is to determine the extent to which these results hold in other visual perceptual tasks and in other sensory modalities, and to explore the origins of this pre-decisional choice-related activity.

## Methods and Materials

### Task and visual stimuli

All procedures were approved by the University of Texas Institutional Animal Care and Use Committee and conformed to NIH standards. Similar task and VSDI processing methods have been described previously (***Chen et al., 2006***, ***2008***). Briefly, monkeys were trained to perform a reaction-time visual detection task. Each trial began after the monkey established fixation on a 0.1° square fixation point displayed against a uniform gray background. The target was a small Gabor patch (sinusoidal grating modulated by a 2-D Gaussian envelope; *σ* = 0.17°, spatial frequency = 2.76 cycles/°, phase = 90°) that appeared at a fixed location (eccentricity = 2.48° for Monkey T, 1.58° for Monkey C). The Gabor stimuli appeared at one or two discrete target contrast levels at or near the monkey’s psychophysical detection threshold (2.6%-5%), and were held constant throughout each session.

During the stimulus-detection task, in order to indicate a ‘target-present’ choice, the monkey was required to shift gaze to the location of the target within 600 ms from target onset time (but not sooner than 75 ms after target onset time) and maintain gaze at that location for an additional 300 ms in order to receive a liquid reward. In target-absent trials, the monkey was required to maintain fixation within a small window (< 1° full width) around the fixation point for an additional 900 ms from target onset time in order to obtain a reward. The target remained on for 300 ms or until the monkey initiated a saccade. On all trials in which the monkey made a saccadic eye movement towards the target location during the relevant temporal interval, a small point appeared at the target location to help the monkey maintain post-saccadic fixation and to provide a visual reminder of the correct target location. The monkeys made a mixture of correct and incorrect responses on both target-absent and target-present trials, allowing us to examine neural activity during Hit, Miss, FA, and CR outcomes (for details see Appendix 1 Table 1). Visual stimuli were presented on a gamma-corrected 21” color display at a fixed mean luminance of 30 cd/m^2^. The display subtended 20.5° x 15.4° at a viewing distance of 108 cm, had a pixel resolution of 1024 x 768, 30-bit color depth, and a refresh rate of 100Hz.

On each day of data collection, the monkey initially performed the detection task in a short block of trials that included a range of target contrasts. One or two contrasts producing a mix of behavioral outcomes were chosen for use in the subsequent imaging experiment.

### VSDI recording and signal pre-processing

While the monkey performed the task, we recorded neural population signals in V1 using the voltage-sensitive dyes RH-1838 or RH-1691 (***Shoham et al., 1999***). Imaging data were obtained using the Imager 3001 system (Optical Imaging, Inc.), collected at a resolution of 512×512 pixels at a sampling rate of 100 Hz.

During analysis, the data were further binned to a resolution of 64×64 pixels, where each pixel corresponds to 0.5×0.5 mm^2^ of cortex. Our basic VSDI signal pre-processing consists of two steps: First, we normalized the responses at each site (a binned group of pixels) by the average fluorescence at that site across all trials and camera frames. This normalization serves to minimize the effects of uneven illumination and staining. Second, we subtracted from each waveform the temporal average over an 80 ms interval centered on the stimulus onset time. This step serves to minimize effects of low temporal frequency fluctuations.

To obtain activity maps, time courses, and pooled scalar responses, we performed several additional steps. To obtain space-averaged time courses (Fig. 8; Fig. 8 – Supplementary Fig. 1) we averaged responses over a rectangular area of 2.5×2.5 mm^2^ centered on the location with the most reliable response (measured as maximal *d*′) obtained on control trials at high (25%) target contrast. More concretely, this *d*′ measure for a candidate pixel is computed as the mean response difference at that pixel between the 25% high contrast and 0% no contrast trials, minus the standard deviation over all trials: *d*′ = (*μ*_25%_ − *μ*_0%_)/*σ*. This is intended to capture dynamics in an aggregate region of interest, and not to represent a plausible pooled response/decision variable as we consider elsewhere in the paper.

To obtain time-averaged spatial activity maps (Fig. 2; Fig. 2 – Supplementary Fig. 1) we averaged responses over a post-stimulus temporal interval beginning 100 ms after stimulus onset and ending at the median response time, or 20 ms before the response time in that particular trial, whichever was earlier. The median RT was 346 ms in Monkey T and 263 ms in Monkey C. We explored other temporal intervals over which to average activity, including intervals that were longer as well as shorter than that used here, and also explored temporal intervals that were time-locked to the saccadic eye movements, but did not find any significant differences. This observation is likely due to low frequency temporal correlations present in the neural population response (***Chen et al., 2008***).

To obtain time-space-averaged scalar responses representing the neural activity on each trial (Fig. 2C; Fig. 2 – Supplementary Fig. 1B,E), we averaged over both space and time in accordance with both of the above procedures. For display purposes (e.g., Fig. 2), we normalized average pooled responses to a standardized ΔF/F by subtracting the mean response during target-absent trials, and dividing the result by the mean response during target-present trials. The activity evoked by our at-threshold stimuli reached approximately 8% of the response to a 25% contrast stimulus. Time-space-averaged scalar responses using different pooling rules (e.g. in the scalar likelihood model, Table 1, Figs. 5, 6) are discussed in the next subsection.

To collate results across multiple data sets, we fit the average stimulus-evoked response to a high contrast (25%) Gabor stimulus (collected during a control block immediately prior to each detection block) with a 2-dimensional Gaussian. We then spatially shifted the responses of each experiment such that the peak of each experiment’s fitted response occurred at a common pixel.

### Pooling algorithms

To explore the impact of various spatial pooling models on our results, we repeated our analysis for a diverse family of 9 alternative pooling rules that each perform a more principled weighted averaging of the VSDI activity maps. To begin, we time-averaged each trial’s responses at each location over the temporal averaging window described above. Then, we normalized the resulting maps analogously to the ΔF/F above by subtracting the mean target-absent map and dividing by the estimated [scalar] peak amplitude of the mean target-present response. This normalization procedure was done on an experiment-by-experiment and a contrast-by-contrast basis to minimize any influence of variation in contrast or experiment quality on the spatial pooling rules.

We then pooled responses over space by the application of various pooling algorithms, with results summarized in Table 1. Each rule is defined by a map of weights, normalized to sum to 1, and the pooled response on each trial was the dot product of that trial’s activity map with the pooling weights:

1. The choice-triggered pooling rule had weights proportional to the 2-D Gaussian that best fit the average response difference between trials ending with a ‘Yes’ choice, and those ending with a ‘No’ choice; to assure this difference is strictly choice-triggered, we preprocess ‘stimulus present’ trials by subtracting the average ‘stimulus present’ response.

2. The stimulus-triggered pooling rule had weights proportional to the 2-D Gaussian that best fit the average response difference between trials with ‘stimulus present’, and those with ‘stimulus absent’.

3–7. Five Gaussian pooling rules had weights proportional to 2-dimensional Gaussians with *σ* = (0.5, 1, 2, 3, 4) mm respectively. This range of parameters spanned the breadth of the 2-D Gaussian that best fit the average stimulus-evoked response maps, which had parameters *σ*_*major*_ = 2.00 mm, *σ*_*minor*_ = 1.71 mm.

8–9. Previous work showed that the optimal pooling rules for pooling decision variables from VSDI activity are difference-of-Gaussian (DOG) weight maps (***Chen et al., 2006***). The last two pooling rules were an optimal choice-triggered pooling rule and optimal stimulus-triggered pooling rule, now with weights proportional to the 2-D DOGs that best fit the respective average response difference described in rules 1 and 2. In practice, these best-fit DOGs have positive weights in the center and negative weights in the surround, which serves to filter the broad spatial correlations observed in VSDI data (***Chen et al., 2006***).

### Sub-sampling Analysis

To examine the impact of our assumption that the subject and researcher share equal access to the sensory population (Fig. 4), we extended our Monte Carlo simulation to two dimensions. The stimulus-evoked response was a two-dimensional Gaussian with peak amplitude of unity that matched the shape of the stimulus-evoked response in the real experiments. As in our real experiments, the stimulus appeared on 50% of trials, and was added to spatially correlated Gaussian noise that occurred on all trials. The spatial noise spectrum was matched to that measured in real experiments. For each of the pooling models we considered, the simulated subject and researcher each applied the model to these sensory signals, producing distributions of pooled sensory response amplitudes. We normalized all pairs of distributions such that the target-absent distribution had a mean of zero, and the target-present distribution had a mean of one. We then added independent zero-mean Gaussian noise to the subject’s pooled sensory responses and independent zero-mean Gaussian noise to the researcher’s pooled sensory responses as in our earlier analysis.

To explore the possibility of unequal access to V1 populations, we performed simulated experiments in which the researcher, the subject, or both, are given independent access only to a random sub-population of the simulated cortex. We then observed the relationships between the estimated values of MF and DF under the model that assumed equal access (Fig. 4), and the actual values that were used to generate the simulation.

Consider the case when the simulated experimenter is given access to 100% of the available cortical signals, while the simulated monkey samples a random subset, consisting of 70%, 80%, 90%, or 100% of the pixels (Fig. 6A-B). Intuitively, as the number of noisy neurons/pixels contributing to the monkey’s pooled response decreases, the monkey’s signal-to-noise ratio decreases. Because we normalize the pooled responses such that the mean target-absent response is equal to zero and the mean target-present response is equal to unity, sub-sampling causes the monkey’s pooled variability to increase. While this additional variability is inherited from the noisy neurons/pixels in our population and thus originates in sensory cortex, our model (which assumes that all noise in sensory cortex is shared; Fig. 4) considers any choice-related variability that is not measured as downstream noise. Thus when the monkey sub-samples the sensory population, we expect our model to overestimate the amount of downstream noise, or in other words, underestimate DF. Fig. 6A shows the relationship between the maximum likelihood estimate of DF and the actual DF value used to generate the simulated experiment. The black line represents the case in which both the simulated experimenter and monkey each sample 100% of the available neurons, and thus our assumption of equal access is met. The grey lines represent cases in which the simulated monkey sub-samples the population. This figure shows that as the monkey sub-samples the population, the model systematically underestimates the true DF, consistent with our intuition.

In contrast to its effect on DF, sub-sampling by the monkey has no effect on MF. Sub-sampling by the monkey should affect neither the simulated experimenter’s pooled sensory signals, nor the total measured variability, and thus the ratio MF should remain unchanged. Fig. 6B, which shows the relationship between the maximum likelihood estimate of MF and the actual MF value used to generate the simulated experiment, verifies this intuition. Cases in which the monkey sub-samples the population are not distinguishable from the case in which the monkey samples 100% of the available cortical signals.

Next consider the case when the simulated monkey is given access to 100% of the available cortical signals, while the simulated experimenter samples a random subset, consisting of 70%, 80%, 90%, or 100% of the pixels (Fig. 6C-D). We expect these cases to show symmetrical results to those described above. On one hand, sub-sampling by the experimenter has no effect on the model’s estimate of DF (Fig. 6C), since this sub-sampling affects neither the monkey’s pooled signals nor the total decision-related variability. However, as the experimenter samples fewer pixels, the experimenter’s SNR decreases, and as above, this loss in SNR manifests as additional noise. While this noise is sensory in origin, our model considers it as measurement noise. Thus in the case when the experimenter sub-samples the population, we expect our model to overestimate measurement noise, or in other words, underestimate MF. Fig. 6D illustrates that in these cases, the model systematically underestimates MF. This could be one reason for the low MF values reported here.

Finally, we considered cases in which both the simulated experimenter and monkey each sample a subset of the sensory cortical signals (Fig. 6E-F). Because sub-sampling by the experimenter has no effect on DF, differences between the estimated and actual DF values in these cases should be related to only the monkey’s sub-sampling. Fig. 6E shows the relationship between the actual and estimated DF when the experimenter samples just 50% of the available neurons and while the monkey samples 70%, 80%, 90%, or 100% of the pixels. This plot is identical to that of Fig. 6A, consistent with our intuition. Similarly, we expect that differences between actual and estimated MF should be driven solely by the degree of sub-sampling by the experimenter. Fig. 6F shows the relationship between the actual and estimated MF when the monkey samples just 50% of the available neurons and while the experimenter samples 70%, 80%, 90%, or 100% of the pixels. This plot is identical to that of Fig. 6B, also consistent with our intuition. We repeated this analysis for seven of our nine pooling models (the two pooling models based on the real monkey’s choices are undefined in the simulation). Results for the stimulus-triggered pooling model are shown in Fig. 6, though our choice of pooling model did not significantly affect our results.

### Eye Movements

To test whether small differences in eye movements may have been responsible for some of our results, we examined two eye movement statistics, which summarized the quality of fixation (1) within each trial, and (2) across trials. The first statistic was the standard deviation of the eye position over the same time interval used for averaging of VSDI signals, and had a mean of 0.0549 degrees and a standard deviation of 0.0121 degrees. The second statistic was the average distance from the monkey’s direction of gaze to the fixation point during this same time interval, and had a mean of 0.0935 degrees and standard deviation of 0.0593 across trials. For each metric, we separated the data into two halves (above median and below median) and examined the population activity maps obtained separately from each half of the data. For each half, we obtained bootstrapped sampling distributions of the VSDI response amplitude averaged over a 2.5×2.5 mm^2^ region centered on the peak of the 2-D Gaussian that best fit the stimulus evoked response to a high contrast (25%) Gabor stimulus collected during control blocks prior to each session. We repeated this analysis separately for Hit and FA trials. We also obtained the probability that the monkey makes each behavioral choice (Hit and FA) for each half of the trials. For both metrics, the half of trials with the poorest fixation quality was associated with an increase in the probability of a FA (an increase of approximately 3%). However, none of the differences in the VSDI amplitude’s mean or standard deviation were significant at the 0.05 level, suggesting that small differences in eye movements are not responsible for our results (see Appendix 1 Table 2). Likewise, repeating the maximum likelihood procedure for the model estimates of MF and DF, under best- and worst-half splits of the data, do not yield significant differences. A significant fraction of the variability in measured eye movements is likely due to variability introduced by the eye tracking device.

### Statistical testing

All statistical tests on the behavioral data, physiological data, and Monte Carlo data were performed using bootstrap methods (***Efron and Tibshirani, 1994***).

### Theoretical models

In order to understand the relationship between measurements of population neural activity in primary visual cortex and behavioral choice, we developed a simple two-channel signal detection model, outlined in Fig. 4. Detailed description of our model derivations and maximum likelihood calculation is given in Supplemental Information. Briefly, we start with the assumption of three independent Gaussian, additive noise sources: sensory noise *n*_*s*_; downstream decision-related noise *n*_*d*_; and decision-unrelated noise affecting the VSDI measurements *n*_*m*_. We have previously shown that VSDI signals are consistent with Gaussian additive noise (***Chen et al., 2006***) and we performed similar tests to verify these properties in our present data (recall Fig. 2E-J). We assume these noise sources to be zero-mean Gaussians with variances 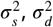, and 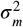, respectively. The subject’s internal decision variable is *r*_*d*_ = *X* + *n*_*s*_ + *n*_*d*_, where *X* = {0, *c*} represents the strength of the visual stimulus response and takes on the value 0 when the target is absent, or the value *c* when the target is present. Because the target was limited to a narrow range of contrasts (2.6%-5.0%), we fit a line to the aggregate data across all threshold target-present contrast levels and all experiments to determine how *c* depends on target contrast. We model the pooled response measured by the experimenters as *r*_*m*_ = *κ*(*X* + *n*_*s*_) + *n*_*m*_ where *κ* represents the dye sensitivity, which determines how strongly V1 activity modulates the VSDI signal and was estimated based on the average amplitude of the response to 25% contrast targets in a control block immediately prior to each detection block. Dye sensitivity varies across experiments due to fluctuations in the quality of staining, so *κ* varies to remove this source of extraneous variability from our model. The subject’s binary decision *D* = {0, 1} represents whether or not the subject reported seeing the target, and is determined by a threshold, *D* = *H* (*r*_*d*_ − *τ*) where *H* (·) is the Heaviside step function and *τ* is the decision criterion. Given these ingredients, the log probability of the subject’s decision *D* and the observed responses *r*_*m*_ on a single trial is given by:

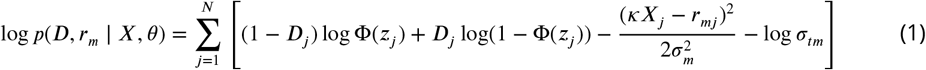

plus a constant that does not depend on the model parameters, where

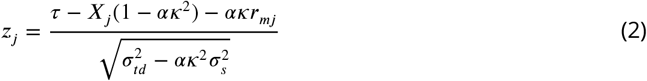

and where Φ(·) is the standard normal cumulative density function (cdf), 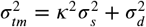 is the total measurement noise variance, 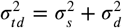 is the total decision noise variance, and 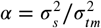 is the ratio of shared to total measurement noise variance. We obtained maximum likelihood fits of the model parameters 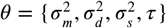 by numerical optimization of the log-likelihood (given by Eq. 1, summed over all trials) using MATLAB’s fmincon.

While DF and MF values were obtained from parameters estimated by the full model (Eq. 1) that takes into account variations in contrast and experiment-to-experiment fluctuations in signal quality, such variations lead to a broadening of the distributions of scalar responses tied to each behavioral category. To visualize the different qualitative modes of our model (Fig. 5), these data-dependent parameters can be fixed to create a simplified model. The simplified model uses a single contrast value (*X* = 1 “On” or *X* = 0 “Off”) with no variations in VSDI signal quality (*κ* = 1). In the simplified model, the average FA and M response difference 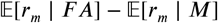 has a linear relationship with DF once the total decision-related variability is fixed (Fig. 5E):

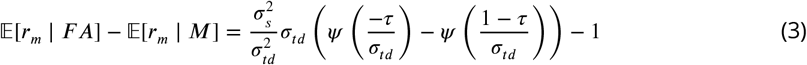

where the leading ratio is exactly DF, and *ψ* (·) = *ϕ*(·)/Φ(·) is the ratio of standard normal pdf to cdf. The −*τ* and 1 − *τ* come from *X* − *τ* when *X* = 0 on FA trials and *X* = 1 on Miss trials. For completeness, we note that the unsimplified version of this equation can be computed using (eq. 16) in the Supplemental Materials.

In order to understand the different predictions for our VSDI data under additive signal noise (in the previous model) versus multiplicative signal gains (an alternate hypothesis), we expanded this scalar likelihood model to a full matrix-valued model that simulates the neural responses at the original pixel resolution of the observed VSDI signals. As in the scalar likelihood model, detailed description of these model derivations is given in Supplemental Information. Briefly, starting from the same scalar model backbone, we define the additive model by assuming the scalar measurement *r*_*m*_ is instead a *p*^2^ × 1 vector *R*_*m*_ (a “flattened” match to the *p* × *p* pixel maps of our VSDI recordings) with *R*_*m*_ = *κ*(*X* + *N*_*s*_) + *N*_*m*_. At the decision stage, we explicitly pool the sensory response with a pooling rule *W*, such that 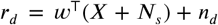. Each of the nine pooling rules we consider (cf. Table 1) correspond to specific choices of *W*. In this model, *N*_*s*_ and *N*_*m*_ become independent multivariate Gaussian additive noise sources with mean 0 and distance-dependent spatial covariances defined by a kernel, and *n*_*d*_ is the same as in the original model.

Analogously, we define the multiplicative model by exchanging the additive *N*_*s*_ for a multiplicative *G*_*s*_, such that *R*_*M*_ = *κ*(*G*_*s*_ ⊙ *X*) + *N*_*m*_ and 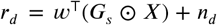, where ⊙ denotes an elementwise (pixelwise) product. In this model, *G*_*s*_ is a matrix in which each pixel is a Gamma distribution with mean 1 and distance-dependent spatial covariances between pixels are generated using the multivariate Gaussian copula. In this way, both models feature spatial correlations noise in measurement and sensory noise. For any pooling rule amongst the 9 used in the scalar likelihood model, we can repurpose the optimized variance fractions of these noise sources given by MF and DF, such that these matrix models would need to optimize only (1) the scale of the spatial correlations to match the VSDI data and (2) the total variance of the model to match the performance (Hits, Misses, FAs, CRs). Noting that the multiplicative model does not admit a closed-form function for the likelihood of *D*, *R*_*m*_ | *X*, the spatial correlation scale and total variance in the multiplicative model were instead optimized by simple grid search over parameters, and in the additive model analytically using metrics from classical signal detection theory. Results from these models are summarized in Figure 7.

#### Applicability of our model to other binary decision tasks

Our model and metric are not restricted to “yes/no” detection tasks but generalize to other types of binary decision tasks. For example, consider the relationship between our detection task and the ubiquitous motion direction discrimination task (e.g. ***Newsome et al., 1989***). In each case, the subject must make a binary decision about a sensory stimulus while responses are collected from neurons that are sensitive to the stimulus presented. In direction discrimination, the subject is presented with a field of moving dots. A fraction of the dots move coherently in one direction while the remaining dots exhibit random motion. The fraction of dots that move coherently is the “coherence”, where lower coherence trials are more difficult to discriminate. Consider a case in which an experimenter identifies a population of neurons with a preferred direction of motion, and makes repeated measurements of their responses to a near-threshold low-coherence stimulus moving in the preferred direction (“preferred”; e.g., leftward motion) and in the opposite (“null”; e.g., rightward motion) direction; the subject must choose “leftward” or “rightward” in an attempt to correctly indicate the direction of motion presented. The collection of pooled responses to a single coherence and two different directions forms a pair of distributions, analogous to our pair of neural response distributions for target-present and target-absent trials, respectively. The experimenter may then normalize these distributions by subtracting from all responses the mean of the responses belonging to the “null” stimulus distribution, then dividing all responses by the mean of the responses belonging to the “preferred” stimulus distribution.

While direction discrimination is a one-interval 2-alternative forced choice task yielding just two possible outcomes (correct and incorrect choices), one may nevertheless identify two distinct distributions of pooled neural responses belonging to incorrect choices. The first distribution is the collection of pooled neural responses to stimuli in the “null” direction in trials in which the subject chose “preferred” (analogous to our FA trials), and the second distribution is the collection of pooled neural responses to stimuli in the “preferred” direction in trials in which the subject chose “null” (analogous to our Miss trials). The equivalent of 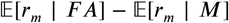 in this task can be defined as the difference between the mean of the former distribution and the mean of the latter distribution. As in our experiments, the difference between these means quantifies the extent to which responses from this neural population correspond more closely, on average, to the stimulus or to the choice. Under the assumption that these pooled neural responses come from Gaussian, equal variance distributions, the experimenter can use maximum likelihood to estimate the same model parameters 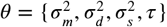 (see Eq. 1).

## Data and Code Availability

Preprocessed data and MATLAB code to reproduce analyses, figures, tables, and simulations reported in this paper are available here: https://bitbucket.org/pillowlab/v1choicevariability

## Acknowledgments

We thank Tihomir Cakic and other members of the Seidemann laboratory for their contributions to this project. This work was supported by NIH/NEI RO1EY016454 and RO1EY16752 (ES), an NSF Graduate Student Fellowship (MJM), and a Ruth L. Kirschstein NRSA (CAM).

## Appendix 1 Supplementary figures and tables

**Appendix 1 Table 1.** Summary of behavioral performance. “Combined” data corresponds to data shown in Fig. 2.

|  | # Experiments | # Hit | # Miss | # FA | # CR | # Total | % Correct | Median RT |
| --- | --- | --- | --- | --- | --- | --- | --- | --- |
| Monkey T | 22 | 2511 | 1303 | 432 | 3341 | 7587 | 77% | 346 |
| Monkey C | 5 | 588 | 238 | 207 | 587 | 1620 | 73% | 263 |
| Combined | 27 | 3099 | 1541 | 639 | 3928 | 9207 | 76% | 334 |

**Appendix 1 Figure 1.**
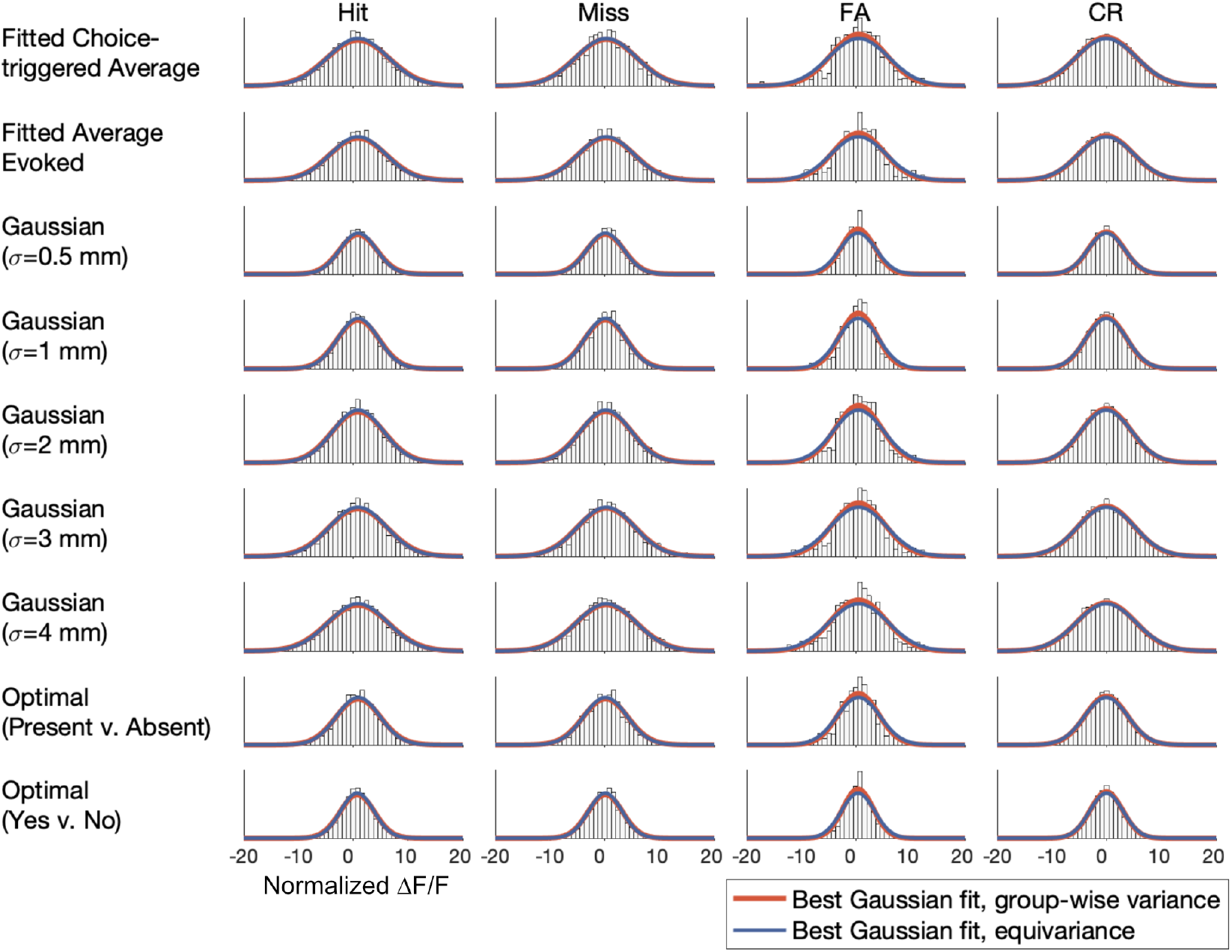
Test of the modeling assumption that the distribution of decision variables (with units ΔF/F) are Gaussian-distributed with equal variance across trial type (Hit, Miss, FA, CR). Each row represents a different pooling rule, and each column a different trial type, and across all combinations we observe that the best-fit Gaussians (thick red lines) are a strong match for the experimental data. Moreover, the best-fit Gaussians with the equivariance constraint (thin blue lines) are nearly indistinguishable from these fits.

**Appendix 1 Table 2.** Eye movement controls. Average VSDI response amplitude and probability of behavioral outcome for Hits (left) and FAs (right), as well as model-fitted measurement and decision variance fractions MF and DF, all reobtained separately for the “best” and “worst” halves of data under two measures of fixation quality (see Methods and Materials). VSDI response amplitudes were averaged over a 2.5×2.5 mm^2^ region centered on the peak location of the 2-D Gaussian that best fit the stimulus-evoked response to a high contrast (25%) Gabor stimulus (collected during control blocks before each imaging session). While the half of trials with the poorest fixation quality was associated with an increase in the probability of a FA (an increase of approximately 3%), none of the differences in average VSDI amplitude or VSDI variability were significant at the 0.05 level, suggesting that small differences in eye movements are not responsible for our results. Likewise, in model-fitting, the MF metric is insensitive and the DF metric is sensitive to subsets of trials, but none of the differences between or within MF and DF were significant at the 0.05 level. Values in parentheses indicate 95% confidence intervals from bootstrapping.

| Response<br>Statistic<br>Fixation<br>Metric |  | Hits |  |  | False Alarms |  |  | Model Estimates |  |
| --- | --- | --- | --- | --- | --- | --- | --- | --- | --- |
|  |  | Amplitude<br>(10<sup>-3</sup>)<br>(2.5x2.5 mm<sup>2</sup>) | Std Dev<br>(10<sup>-3</sup>)<br>(2.5x2.5 mm<sup>2</sup>) | probability of Hit | Amplitude<br>(10<sup>-3</sup>)<br>(2.5x2.5 mm<sup>2</sup>) | Std Dev<br>(10<sup>-3</sup>)<br>(2.5x2.5 mm<sup>2</sup>) | probability of FA | MF | DF |
| Within-trial<br>fixation | Best half | 0.46 (0.37, 0.56) | 1.92 (1.83, 2.01) | 0.68 (0.66, 0.70) | 0.27 (0.07, 0.47) | 1.73 (1.57, 1.89) | 0.12 (0.11, 0.14)* | 0.02 (0.01, 0.26) | 0.92 (0.31, 1.00) |
|  | Worst half | 0.46 (0.37, 0.56) | 1.90 (1.81, 1.99) | 0.66 (0.64, 0.68) | 0.25 (0.07, 0.43) | 1.74 (1.58, 1.88) | 0.16 (0.14, 0.17)* | 0.02 (0.00, 0.10) | 0.64 (0.12, 1.00) |
| Across-trial<br>fixation | Best half | 0.53 (0.43, 0.63) | 1.92 (1.84, 2.00) | 0.67 (0.65, 0.69) | 0.13 (-0.07, 0.33) | 1.74 (1.56, 1.90) | 0.12 (0.11, 0.14)* | 0.02 (0.01, 0.06) | 0.62 (0.19, 1.00) |
|  | Worst half | 0.39 (0.30, 0.49) | 1.90 (1.80, 1.99) | 0.67 (0.65, 0.69) | 0.37 (0.19, 0.55) | 1.73 (1.58, 1.87) | 0.16 (0.14, 0.17)* | 0.03 (0.00, 0.09) | 0.93 (0.31, 1.00) |

## Appendix 2 Theoretical models and derivations

## Scalar-valued likelihood model formulation

Suppose we have a stimulus variable *X* that takes on the value 0 when signal is absent, or contrast *c* when signal is present. We will assume that *X* influences two (correlated, possibly overlapping) measures of neural activity: a *measured* neural response *r*_*m*_ and an intrinsic *decision-related* response *r*_*d*_. We will assume that the animal’s decision *D* on a single trial results from thresholding *r*_*d*_,

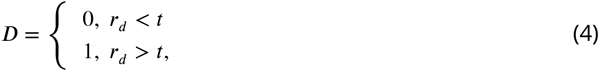

where *t* is a *criterion* for the animal’s decision. In experimental settings, we have access to *X*, *r*_*m*_ and *D* on each trial, but do not observe *r*_*d*_ directly. Below, we will define a model of the statistical relationship between these variables, and derive formulas for several quantities of interest.

We assume three independent additive noise sources: (1) “sensory noise” present in V1, which affects *r*_*m*_ and *r*_*d*_; (2) downstream decision noise that affects *r*_*d*_ only; (3) measurement noise that affects *r*_*m*_ only. The model’s behavior is governed by parameters:

- 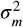 — variance of independent noise added to *r*_*m*_
- 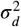 — variance of independent noise added to *r*_*d*_
- 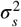 — variance of “shared” noise that is added to *r*_*m*_ and *r*_*d*_.
- *t* — the criterion, or threshold.
- *κ* — the dye sensitivity, determining how strongly V1 activity modulates the VSDI signal.

The model can be described mathematically as follows:

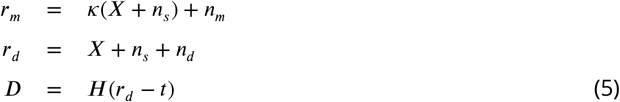

where *n*_*m*_, *n*_*d*_ and *n*_*s*_ are Gaussian random variables with variance 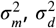 and 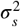, respectively, and *H*(·) is the Heaviside step function.

From this setup, we can derive:

- the total measurement noise: 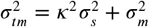
- the total decision noise: 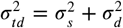

We can write the joint distribution of measurement and decision variables given the stimulus as:

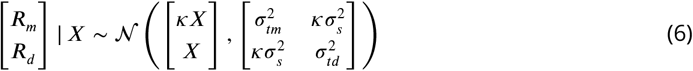

## Deriving the likelihood

The probability of *r*_*m*_ given *X* is given by:

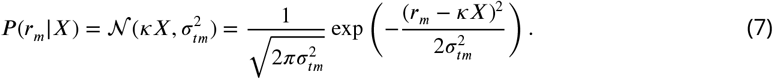

The probability of an “absent” (D=0) or “present” (D=1) response given stimulus *X* alone is the probability that *r*_*d*_ is below or above *t*, which is given by the integrals:

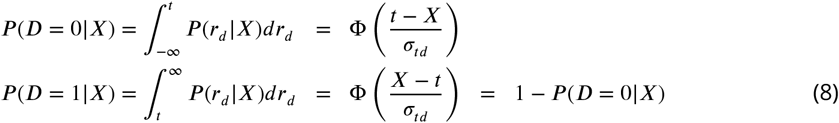

where Φ(·) is the cumulative normal distribution function. These equations will entirely determine the distribution of Hits, False Alarms, Misses and Correct Rejects. (More on this below).

Since *r*_*m*_ and *r*_*d*_ are not independent, we need the distribution of *D* conditioned on *X* and *r*_*m*_ in order to compute the likelihood. From the formula for conditional probabilities of a Gaussian variable, we have:

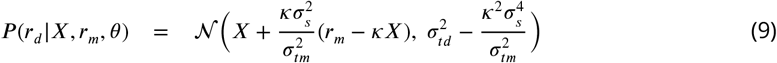

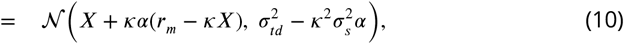

where 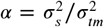 is the fraction of shared noise in the measurement. From this expression, the probability of *D* given *X* and *r*_*m*_ is given by the probability mass under a Gaussian to the left or right of the criterion *t*. Thus, we have

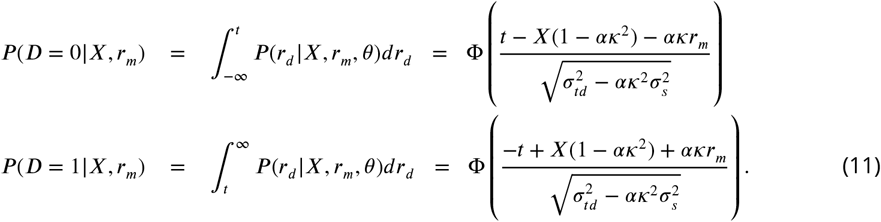

To compute the likelihood function for 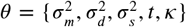, defined as *P*(*D*, *r*_*m*_|*X*, *θ*), we simply take the product *P*(*r*_*m*_|*X*) and *P*(*D*|*X*, *r*_*m*_) as given above, and compute the product over (assumed independent) trials.

We can write the log-likelihood function as follows:

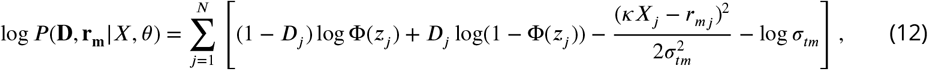

where

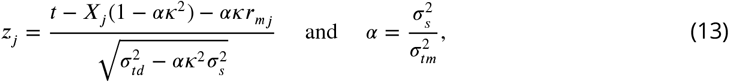

where *j* indexes the trials from 1 to *N*.

Fitting of the model parameters in this formulation can be performed using Matlab’s fmincon to descend the negative of the log-likelihood as a function of the model parameters *θ*.

## Deriving other quantities of interest

In the following, Φ(·) and *ϕ*(·) denote the standard normal cdf and pdf functions, respectively.

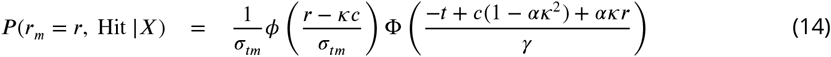

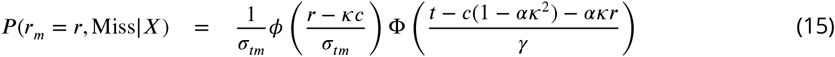

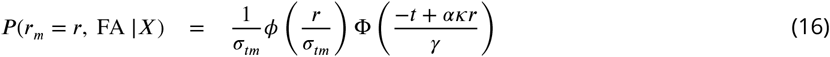

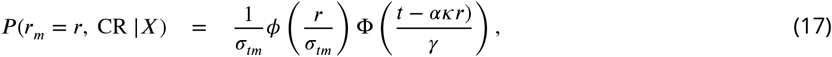

where 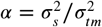 and 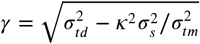, and *c* is the contrast when the stimulus is present.

We can express this more concisely as:

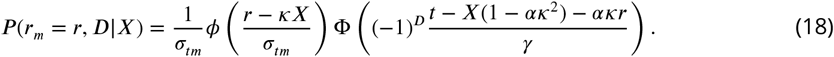

This is the same formula as that of the log-likelihood given above.

If we instead want the conditional distribution over *R*_*m*_ given *X* and *D*, we divide by *P*(*D*|*X*), to obtain:

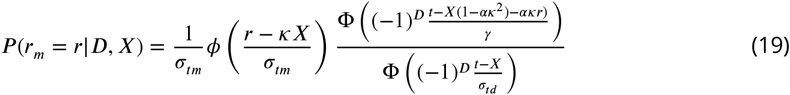

We can define neural sensitivity in terms of d-prime:

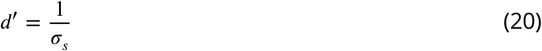

or as area under the ROC curve

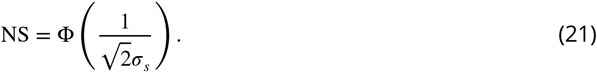

Choice probability can be derived as follows. Let *f*_0_ and *f*_1_ denote the pdfs of *r*_*m*_|*X*, D=0 and *r*_*m*_|*X*, D=1, respectively, which are given by ((19)). Choice probability (CP) is given by

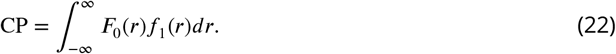

This 1D integral can be computed efficiently using numerical quadrature methods.

## Vector-valued schematic model formulation

Suppose we wanted to extend this scalar-valued likelihood model to vector-valued signals, in order to compare predictions made by additive noise versus multiplicative gain for driving shared [co-]variability in V1 VSDI activity maps. We now assume that the measured neural response *R*_*m*_ is a *p*^2^ × 1 vector, instead of a scalar *r*_*m*_, a “flattened” equivalent of the *p* × *p* grid of pixels in the recorded VSDI images. The same scalar decision-related response *r*_*d*_ is obtained via a linear pooling of the neural response by a *p*^2^ × 1 pooling vector **w**; all nine pooling rules discussed in the scalar model (cf. Table 1) are specific choices of this **w**.

The corresponding signal and measurement noises become vectors *N*_*s*_ and *N*_*m*_ with *covariances* 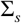 and 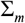, importantly permitting spatial covariability in the VSDI responses. This extended model can be described mathematically in the following *additive noise model* with correlated spatial variability:

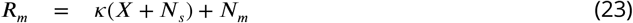

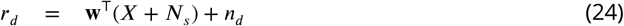

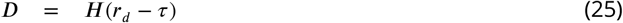

To parametrize these new spatial covariances, assign each pixel of the response *R*_*m*_ with its coordinate location *z* in the image. We can represent the correlating influence of each pixel on other pixels using a kernel *κ*(·, ·): using an exponential kernel, given the broad spatial correlation in our data, the full correlation matrix *κ* is a *p*^2^ × *p*^2^ matrix and the covariance of pixel *i* and *j* given by

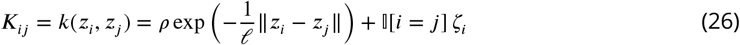

The hyperparameters *ρ* and 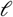 are the magnitude and lengthscale of the spatial correlations, respectively, ∥ · ∥ is the *L*_2_ (Euclidean) norm, and *ζ*_*i*_ are uncorrelated pixelwise variances; in this notation, these independent noise terms are only included if *i* = *j*, represented by the indicator function 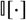. Generating noise with this structure, *N*_*m*_, *N*_*s*_ are multivariate Gaussian random variables with covariances *K*_*m*_ and *K*_*s*_, respectively, and *n*_*d*_ is the same univariate Gaussian random variable with variance 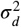. We assume that spatial noises *N*_*s*_ and *N*_*m*_ share a common lengthscale 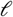, such that our vector-valued model has parameters 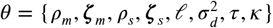.

As an alternate hypothesis, suppose that the *shared* variability is a multiplicative gain, not an additive noise. We still assume the measurement noise is additive, since instrumental noise from the dye and camera are ostensibly additive, and the decision noise is additive to keep these models minimally different. We seek to replace additive signal noise vector *N*_*s*_ with a multiplicative gain vector *G*_*s*_, having mean 1 everywhere such that the mean response to the gain-modulated signal remains the stimulus *X* itself. To inherit spatial covariability featured in the additive noise, we can generate spatially correlated Gamma random variables using the multivariate Gaussian copula. The “copula trick” is a general tool to generate dependent multivariate samples with arbitrary marginal distributions, and is defined in our case by the following sequence of nonlinear transformations:

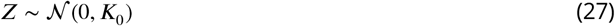

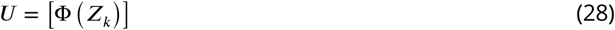

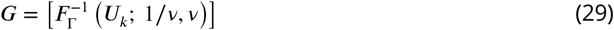

In words, first, we generate zero-mean multivariate normal samples *Z* with *correlation* matrix *K*_0_, which can be achieved with the spatial covariability structure in eq. (26) if we assert *ρ* = 1 and all *ζ* = 0. Next, we use the standard normal CDF Φ(·) to transform these samples elementwise into uniform samples *U* bounded in [0, 1]. Last, these samples can be mapped elementwise onto Gamma distributions using the inverse Gamma CDF 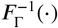; the parametrization in terms of shape 1/*ν* > 0 and scale *ν* > 0 asserts that each pixel in *G* has mean 1 and variance *ν*. Compared to sampling repeatedly from a univariate Gamma distribution Γ(1/*ν*, *ν*) to populate the matrix *G*, these samples are dependent, inheriting the covariability in *K*_0_ into higher-order moments.

Modifying the derivation appropriately yields the *multiplicative gain model* with correlated spatial variability:

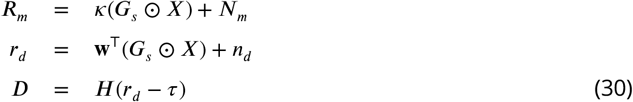

where ⊙ is the Hadamard elementwise product, *N*_*m*_ and *N*_*d*_ are as defined in the additive noise model, and *G*_*s*_ are mean-1 Gamma-distributed gains generated from the copula trick above. Likewise assuming a spatial lengthscale shared amongst all variables, this model has parameters *θ* = 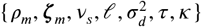. Since the copula transformations are all nonlinear, the correlations between gains in *G*_*s*_ will not be equal to the correlations between their parent samples *Z*. For our goals with these simulations, though, their qualitative similarity is sufficient.

Problematically, the vector-valued multiplicative gain model does not have a well-defined likelihood function for optimization; copula trick notwithstanding, the distribution of the sum of Gamma and Gaussian random variables does not have a useful form. Recalling that our goals with these simulations is only to qualitatively affirm that the VSDI responses are most consistent with additive versus multiplicative noise, a schematic-level optimization is sufficient and tractable if we repurpose insights from the scalar model.

For any pooling rule amongst the 9 used in the scalar likelihood model, we can repurpose the optimized variance fractions of the three noise sources (signal, measurement, decision) given by MF and DF, such that these matrix models would need to optimize only (1) the scale of the spatial correlations to match the VSDI data and (2) the total variance of the model to match the performance (Hits, Misses, FAs, CRs).

## Optimizing the vector-valued simulations

To begin, we define the vector-valued stimulus *X* as the best-fit Gaussian to the average VSDI response on target-present trials. To simplify, we assert *κ* = 1, renormalize *X* such that **w**^T^*X* = 1, and choose the Gaussian pooling rule with *σ* = 2.0mm (pooling rule #5) without loss of generality. We define the spatial lengthscale 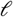 and magnitudes *ρ* as those that minimize (by simple gridsearch) the MSE between the VSDI autocorrelogram on target-present trials and the resulting kernel 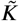 of the form in eq. (26); we use the residual pixelwise variances not captured by this kernel to define ***ζ***.

We also fix the variance fractions DF = 0.86 and MF = 0.04 (cf. Table 1) to their optimal values under that pooling rule, to constrain the [co-]variances such that the only effective degree of freedom remaining is the *total variance* amongst them all. To connect the vector-valued models to the scalar likelihood model, we can pool the responses *r*_*m*_ = **w**^T^*R*_*m*_ to recover the equivalent variances 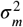 and 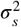 used in the scalar model. Starting with the additive model, we can show that all of *ρ*_*m*_, *ρ*_*s*_, 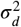 can be expressed analytically in terms of 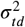. MF and DF are defined in closed form as

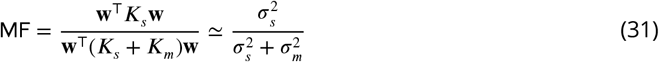

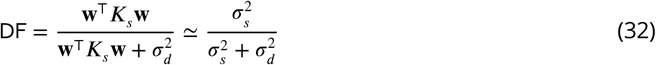

In the additive Gaussian case, the animals’ performance levels, in terms of their Hit and FA rates *H* and *FA*, respectively, imply a certain noise level in the decision variable *r*_*d*_ via its d-prime statistic of *r*_*d*_ | *X*:

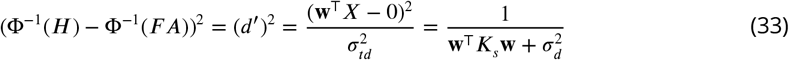

Working backwards with eqs. (31) and (32) to reach the kernel 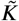, we can define each of the following to generate an additive noise model that robustly matches the observed data:

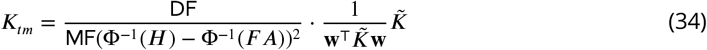

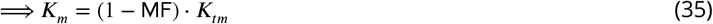

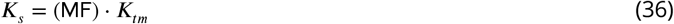

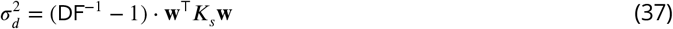

If we simulate a full dataset according to eqs. (23) with these variances and covariances, there exists a decision threshold *τ* that perfectly generates the observed performances *H* and *FA*.

In the multiplicative Gamma case, this procedure is not possible in closed form, as the distributions *r*_*d*_ | *X* are not Gaussian. Moreover, MF and DF are only defined conditioned on the stimulus *X*:

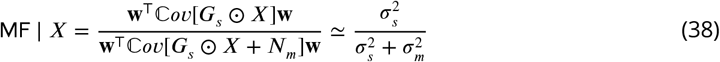

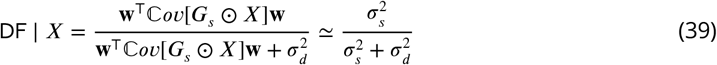

Instead, we can gridsearch over the two effective parameters remaining: the Gamma scale *ν* and a normalizer for *K*_*tm*_ which we will call *β* (cf. eq. (34)). We can loop through the following:

1. (Outer loop) Propose *β* to define 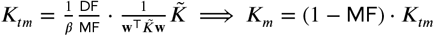. This defines *N*_*m*_.
2. (Inner loop) Adjust *ν* and simulate *G*_*s*_ until eq. (38) is satisfied. This defines *G*_*s*_.
3. (Simulate) Define 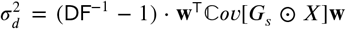 to complete, and simulate full dataset according to eqs. (30).
4. (Optimize) Select decision threshold *τ* that yields performances matching the observed *H* and *FA* (minimizes MSE distance). *If performance is too weak, decrease β and repeat. If performance is too strong, increase β and repeat. Given the constraints, the solution is unique.*

Both the analytical procedure for the additive noise model, and the heuristic procedure for the multiplicative gain model, are lengthy and complex, but both produce schematic simulations that robustly capture qualitative features of the data, including spatial VSDI activity not captured by the pooled likelihood model. Simulating many trials under these two models, we can visualize and compare these data’s predictions to the empirical VSDI data. This is captured in Fig. 7 in the main text.

**Figure 2–Figure supplement 1.**
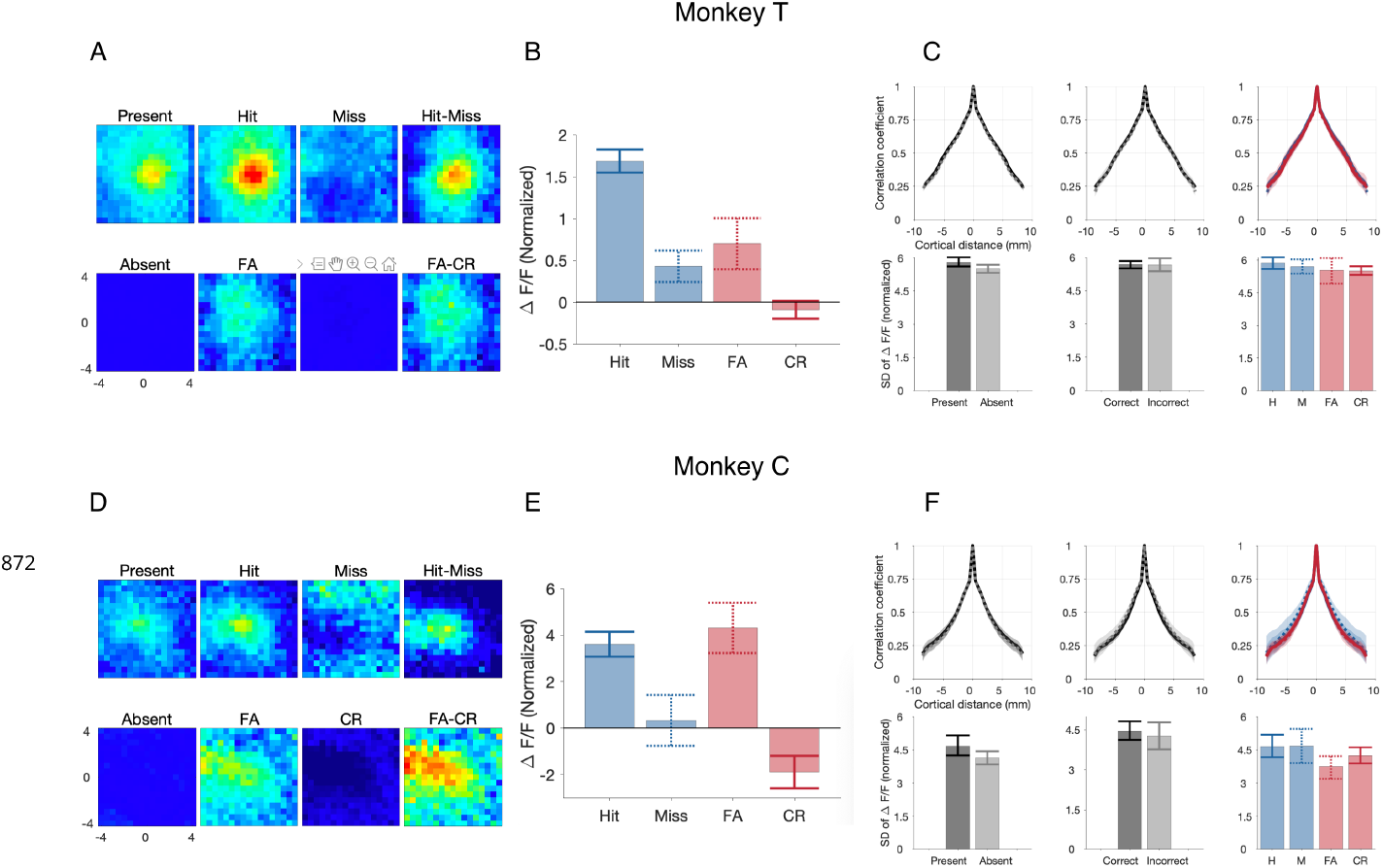
Choice-related population activity in V1, per animal. **A,B,C**, Aggregate summary of VSDI results collapsed over 22 experiments in Monkey T, analogously to Figure 2B,C,D-I, resp. **D,E,F**, Aggregate summary of VSDI results collapsed over 5 experiments in Monkey C, again analogously to Figure 2B,C,D-I, resp.

**Figure 8–Figure supplement 1.**
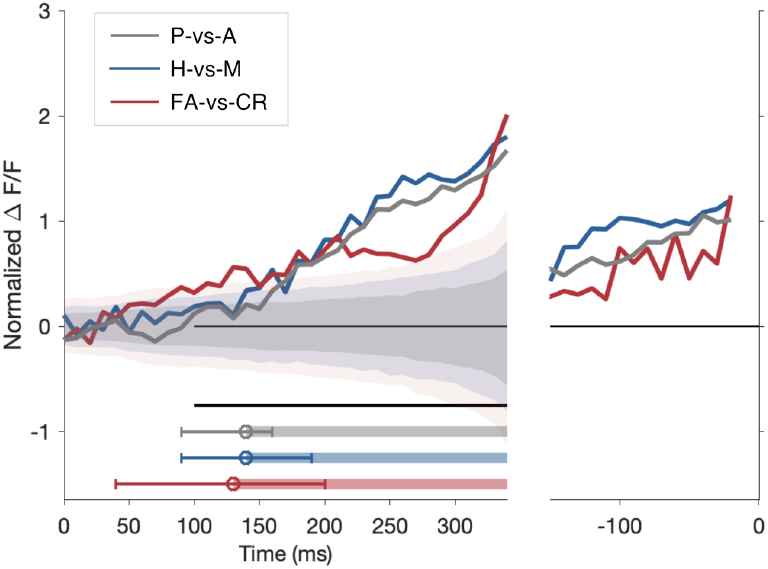
Temporal dynamics of choice-related activity. Time evolution of choice-related differences during target-absent trials (red, mean ‘FA’ minus mean ‘CR’ activity) and target-present trials (blue, mean Hit minus mean Miss activity) are shown alongside the time evolution of the stimulus-evoked response (gray, mean ‘P’ minus mean ‘A’). These lines are the differences of those in the main Fig. 8. Shaded areas in this figure represent 95% bootstrapped confidence intervals on the discriminability of one trial type from the other (Hit vs. Miss, etc.). Significance is marked beneath the curve(s) when the data traces move outside the corresponding shaded intervals.

## Notes

### Competing Interest Statement

The authors have declared no competing interest.

